# Charting developmental trajectories of dynamic brain networks during emotional face processing

**DOI:** 10.64898/2026.04.21.718685

**Authors:** LM Bailey, SC Coleman, N Rhodes, J Crosbie, R Schachar, R Nicolson, E Kelley, J Jones, J Frei, JP Lerch, E Anagnostou, MJ Taylor

## Abstract

Emotional face processing is a critical component in the development of social cognition through childhood. The neural mechanisms supporting this development can be understood by tracking age-related changes in brain activity. Prior work in this area with MEG relies on static, bandlimited, or region-specific measures, which do not capture the dynamic, distributed nature of brain activity. Here we used Dynamic Network Modes (DyNeMo) to analyze MEG data from a large cohort of typically developing individuals (N=224, ages 5-40) during an emotional faces vigilance task. DyNeMo is a data-driven, generative modelling approach which captures functional networks as a set of whole-brain spatiospectral “modes” whose relative mixture (i.e., activation levels) can change dynamically in response to stimuli. We inferred six modes from the MEG data and characterized developmental trajectories in task-related activation and mode connectivity. Across modes we observed distinct developmental trajectories in both measures. With respect to mode activation, a visual mode and a frontotemporal mode (whose labels reflect their respective spatial profiles) increased nonlinearly with age; meanwhile, activation in temporal and sensorimotor modes decreased linearly with age. Meanwhile, connectivity broadly increased with age in all modes, but with different degrees of nonlinearity. These results suggest developmental dissociations between different modes (e.g., visual versus sensorimotor), as well as within individual modes (task-related mode activation versus connectivity). These results provide a comprehensive and complex picture of functional network development underlying emotional face processing.

**Significance Statement:** Brain networks supporting social cognition undergo profound changes from childhood through adolescence to adulthood. However, current understanding of how these networks develop has been limited by conventional analyses of brain imaging data, which typically provide a static picture of brain activity. Here we leveraged a cutting-edge, data-driven modeling approach (DyNeMo) to characterize age-related changes in distributed and dynamic networks supporting emotional face processing, in a large cohort of children and young adults (N=224). We inferred six functional networks whose activation levels changed rapidly in response to emotional faces. Network activation and connectivity exhibited profound and distinct non-linear changes with age, indicating that emotional face processing is supported by complex interactions among multiple dynamic networks, each with different maturational trajectories.

## Introduction

Emotional face processing is fundamental to social cognition and an important component of typical cognitive development. The ability to process faces is not fully developed at birth, but rather matures throughout childhood and adolescence^1–4^. Understanding typical developmental trajectories in emotional face processing, particularly at the neural level, is therefore essential for characterising typical brain and cognitive development.

Much neuroimaging work has sought to characterise the neural correlates of face processing and their typical developmental trajectories. For example, neurophysiological recordings with magneto- and electroencephalography (M/EEG) have consistently revealed developmental shifts in the amplitude, peak latency, and face selectivity of the N/M170 evoked response (typically localized to the fusiform gyri) elicited by images of emotional faces^3,5–8^. Work with functional magnetic resonance imaging (fMRI) has similarly reported increasing face selectivity in the fusiform gyri between early childhood and adulthood^9,10^. Despite this prior emphasis on the fusiform gyrus, more recent work has taken a broader, network-level view of face processing. In this vein, MEG studies have implicated a broad bilateral face processing network comprising occipital, temporal (notably including the fusiform gyri), insular, and orbitofrontal cortices, as well as the anterior cingulate (ACC), and amygdalae^5,11–16^. The functional properties of this network - that is, the magnitude and extent of responses within network nodes (regions), as well as the strength of long-range functional connections between them - mature throughout childhood and adolescence in typically developing (TD) populations^4,7,17–20^. For example, an MEG study by Safar and colleagues^18^ examined frequency-specific connectivity in response to emotional faces, in TDs between the ages of 6 and 40. These authors found developmental increases in connectivity strength in theta and beta frequency bands, with a corresponding decrease in gamma connectivity, between nodes of the canonical face processing network. These findings broadly suggest that cognitive development of emotional face processing is grounded in neural "tuning" of the of the underlying network.

Despite substantial research in this area, efforts to understand network-level developmental trajectories underlying face processing have been hindered by conventional approaches to neuroimaging analysis. *A priori* definition of network hubs for connectivity analyses often varies between studies, depending on which areas are considered relevant to face processing. Moreover, in the context of MEG, connectivity analyses are typically constrained to pre-defined frequency bands and/or time windows of interest (e.g., ^11,16,18,21^). These practices impose two major limitations. First, they are blind to potential effects occurring beyond anticipated regions, frequency bands, and timepoints. Second, they typically provide a temporally static picture of functional networks: for example, by computing connectivity within a pe-defined time window, one loses information about how connectivity might vary over short timescales. More generally, focussing on individual regions or networks ignores the possibility that cognition may be supported by widespread changes across *multiple* networks.

Recent advances in data-driven analysis methods may help circumvent these limitations of conventional hypothesis-driven analyses. In particular, Dynamic Network Modes (DyNeMo)^22,23^ is a generative modelling approach which assumes that observed neural time series data arise from a set of whole-brain *modes* (i.e., functionally distinct network profiles). Each mode has a unique spatiospectral and connectivity profile and, instead of being derived from researcher-defined parameters, they are learned directly from the data. Thus, these methods provide an unbiased means of identifying intrinsic functional networks without reference to hypothesis-driven parameters. DyNeMo allows multiple modes to be active simultaneously, such that the observed data are generated by a linear mixture of modes which may be perturbed by experimental events over short time scales^22–24^. For example, Gohil et al.^23^ demonstrated rapid changes in mode activation levels following voluntary movement and visual stimulus presentation. Critically, mode activation as characterised by DyNeMo captures both localised and distributed neural processes, and therefore provides an elegant bridge between two pillars of human neuroscience: that is, region-specific evoked or induced responses, and transient changes in network-level functional connectivity. As such, DyNeMo provides a parsimonious and data-driven approach for exploring dynamic, task-related network behaviour in neural time series.

In this cross-section ageing study we applied DyNeMo to MEG data acquired during an emotional face processing task, in a large cohort of typically developing participants ranging from 5 to 40 years of age. We capitalized on DyNeMo’s uniquely broad characterization of dynamic network activity to describe developmental trajectories of distributed whole-brain responses and connectivity underlying emotional face processing.

## Results

### Inferred whole-brain modes

We acquired MEG data from a large cohort of typically developing participants (free of neurodevelopmental, psychiatric, or neurological disorders; *N* = 224, ages 5-40) during an emotional face processing task. Participants viewed images of happy and angry faces and responded with a right-handed button press to target trials, on which faces were bordered by a target colour (approximately 25% of trials). We used DyNeMo to train a generative model on preprocessed, spatially-resolved, and time-delay embedded (TDE) continuous MEG data from all participants. We inferred six whole-brain *modes*, each with a unique spatiospectral and connectivity profile (Figure 1).

**Figure 1.**
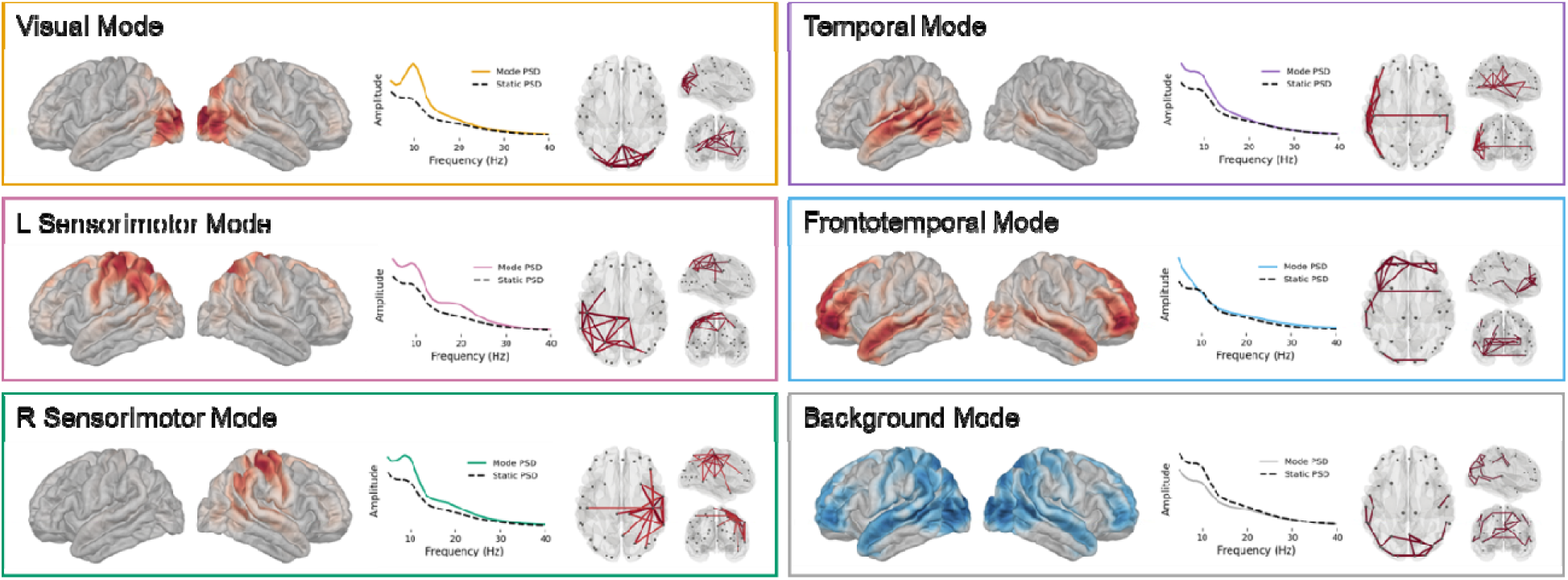
Six modes inferred by DyNeMo. For each mode, spatial maps on brain surfaces show mean power change from the static PSD (power spectral density). Coloured lines are the average mode PSD, dashed lines are the static PSD. Lines overlaid on glass brains are the top 3% of edges from the coherence matrix in the theta to beta range (4-30 Hz).

We observed a visual alpha mode, two sensorimotor alpha/beta modes, a left-sided temporal theta/alpha mode, a frontotemporal theta mode, and a low-power background mode. Note that the spatial map for the frontotemporal mode also encompassed the anterior cingulate (ACC – see S Figure 1 for medial views).

## Task-related mode dynamics

We explored how emotional face processing affected the temporal dynamics (i.e., rapid changes in relative activation) of each mode. Importantly, the generative model used by DyNeMo is trained on continuous data and is blind to experimental events. After model training, we investigated how such events modulated the continuous mode mixture through time. We epoched all mode timecourses relative to the onset of face stimuli during non-target trials. As a secondary verification that DyNeMo was tapping expected task-related modulation of observed modes, we separately epoched the time courses relative to button presses.

Figure 2a shows group-average mode timecourses following emotional face presentation. The visual mode exhibited an early increase in relative activation, consistent with an early visual response, peaking around 125ms post stimulus and followed by a sharp drop-off in activation. This visual deactivation was accompanied by increased activation in other modes, mainly the frontotemporal, right sensorimotor, and temporal modes. Figure 2b shows the mean timecourse of the two sensorimotor modes (which we expected to be modulated by right-handed responses) relative to button presses. Both sensorimotor modes exhibited a clear activation peak immediately following the button press, with the left peak being earlier and lager, consistent with execution of a unilateral motor response.

**Figure 2.**
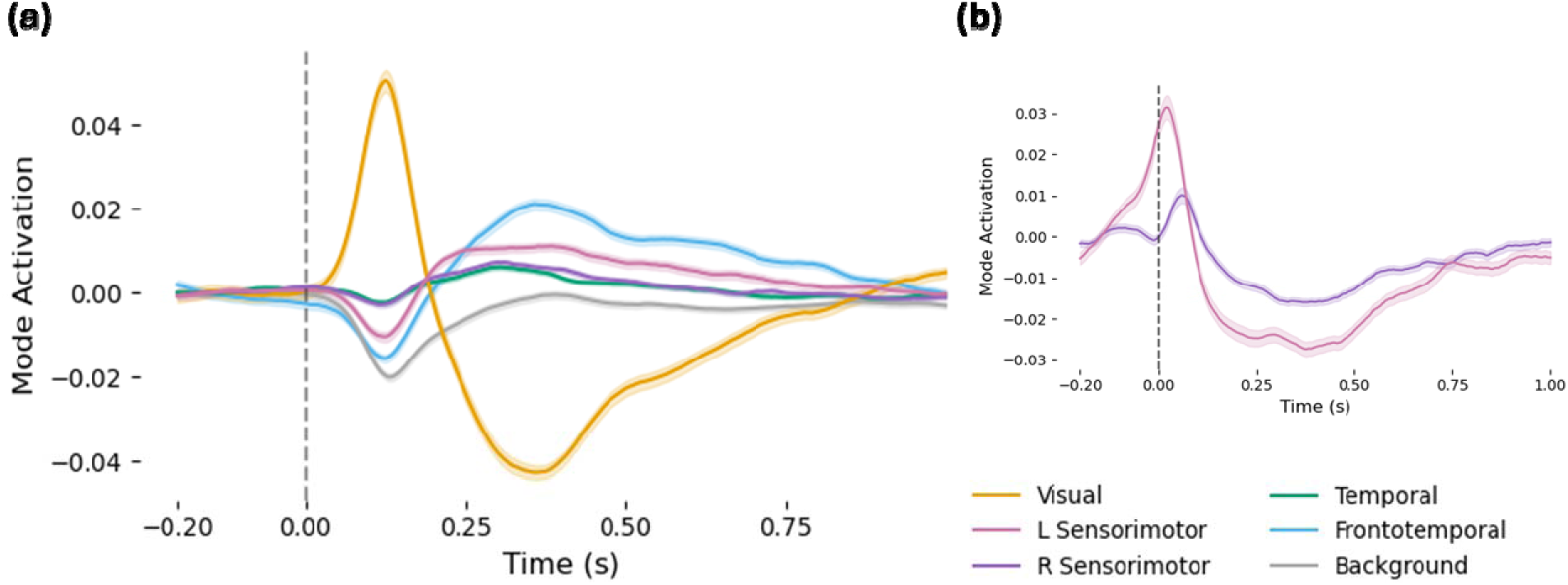
Dynamic task-induced changes in mode activation. **(a)** Mean mode timecourses relative to stimulus presentation (at 0ms, dashed line) on correct non-target trials, averaged across subjects. Shading reflects ±1 SE. **(b)** The mean timecourse of the left sensorimotor mode relative to button presses (0ms, dashed line). All time courses were baseline-corrected for visualisation purposes only.

Given the dynamic nature of mode responses (Figure 2), we conducted a supplementary analysis to investigate which time points were driving the observed age effects; results are presented in Supplementary Materials. In brief, age effects in the visual and frontotemporal modes appeared to be predominantly driven by the activation peaks (∼100ms, 250ms respectively) following stimulus presentation. Age effects in the temporal and right sensorimotor modes were driven by early de-activation (∼100ms), while age effects in the remaining modes were not clearly related to group-level activation peaks.

### Effects of emotion on mode dynamics

We also investigated the extent to which the inferred mode timecourses differed between happy and angry faces. For each time point in the epoched mode timecourses, we conducted a Bayesian t-test between activation values for happy versus angry trials. This yielded a Bayes factor (BF_10_) for each timepoint, per mode. These Bayes factors may be interpreted as a quantitative measure of strength of evidence for a difference in mode activation between happy and angry faces. Mode timecourses for happy and angry faces, and corresponding Bayes factors, are shown in Figure 3. In the visual and frontotemporal modes we saw concurrent between-condition differences at around 600 ms following stimulus presentation. Notably, the evidential strength for this difference was considerably higher in the visual mode (Max BF_10_ ≈ 1600) compared to the frontotemporal mode (Max BF_10_ = 5.89)*. No other modes showed credible evidence for between-condition differences (see S Figure 2 for results in all modes).

**Figure 3.**
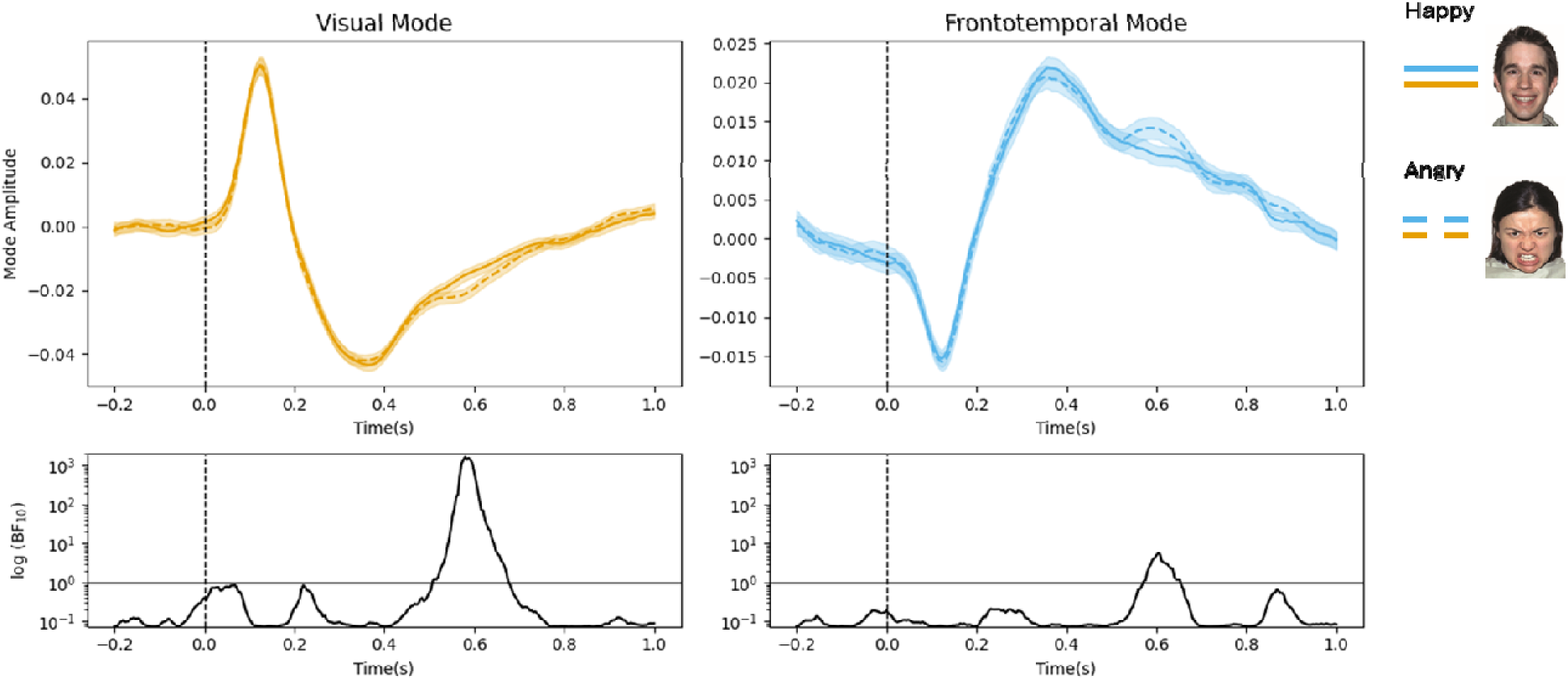
Effect of emotion on mode activation in the visual and frontotemporal modes. **Top row:** Mean mode timecourses relative to the onset of happy (solid lines) and angry (dashed lines) faces. Shading reflects ±1 SE. Timecourses were baseline-corrected for visualisation purposes only. **Bottom row:** Log-transformed Bayes factors (BF_10_) computed for each time point, reflecting the strength of evidence for a difference between happy and angry faces. Solid horizontal lines denote BF_10_ = 1.0, above which values support a difference between conditions. Dashed vertical lines in all panels denote time of stimulus onset.

### Developmental trajectories of mode dynamics

We used Bayesian nonlinear regression to estimate the effect of age on average mode activation in the first 750 ms following face presentation (a full rationale for Bayesian parameter estimation is provided in Methods). The results (Figure 4) indicate different developmental trajectories in each mode. Note that 95% highest-density intervals (HDIs) shown in the right column of Figure 4 reflect the credible range of local slope estimates in each of the seven age bands. Values outside the HDIs can be considered as non-credible with 95% probability (see Methods).

**Figure 4.**
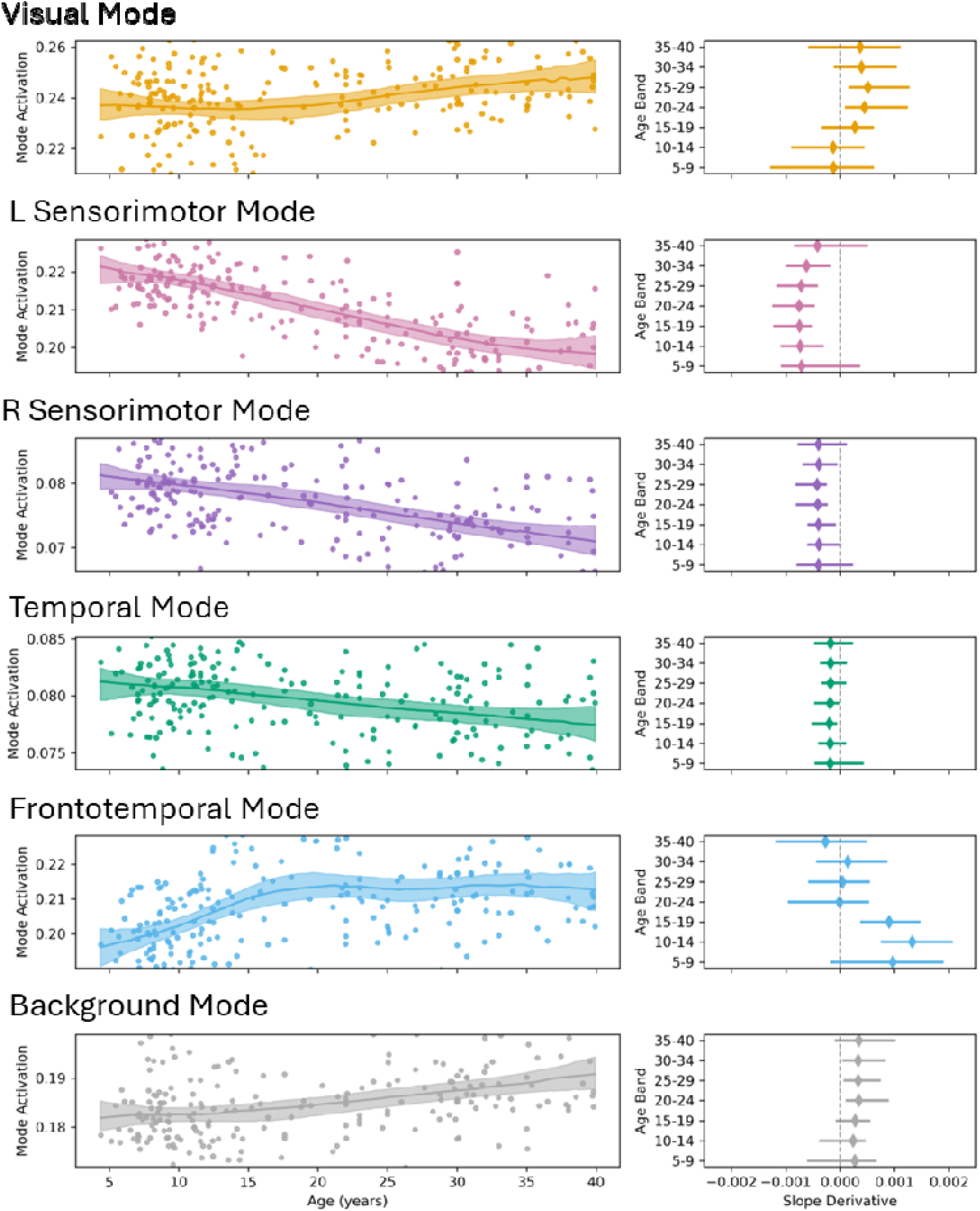
Age-related changes in mode activation in the first 750ms following emotional face presentation. **Left column:** Lines are the most probable regression slopes for the relation between age and mode amplitude; shading shows 95% highest density intervals (HDIs). Points are individual participant means (average across epochs). Points are values for individual participants, averaged across epochs. Outliers above and below the 5^th^ and 95^th^ percentiles, respectively, are not shown. **Right column:** Slope derivatives across age bands. Diamonds denote the most probable estimate, bars denote 95% HDIs.

The frontotemporal mode exhibited the most marked nonlinear effect: activation increased sharply with age until around 20 years of age, after which it plateaued. By contrast, the visual mode exhibited age-related increases mainly throughout early-mid adulthood. Activation in the temporal and sensorimotor modes decreased with age in a roughly linear fashion. We note that the temporal mode exhibited the smallest effect (in terms of magnitude) overall, with HDIs close to or crossing zero in all age bands. Overall, these results indicate that the different modes of neural activity underwent different rates and periods of maturation.

### Developmental trajectories of mode coherence

For each mode, we computed participant-level mean coherence in the top 3% of network edges at the group level (visible in Figures 1 and 5). We again used Bayesian parameter estimation to characterise non-linear relations between age and coherence, for each mode^†^. The left sensorimotor and frontotemporal modes exhibited roughly linear age-related increases in mode coherence, while the visual, right sensorimotor and background modes exhibited more nonlinear effects. In the visual and background modes, age-related changes were maximal during childhood and adolescence (∼5-20 years of age) and then plateaued after ∼30 years of age. In the right sensorimotor and temporal modes, mode coherence appeared to mature later, with maximal changes throughout adolescence and early adulthood.

**Figure 5.**
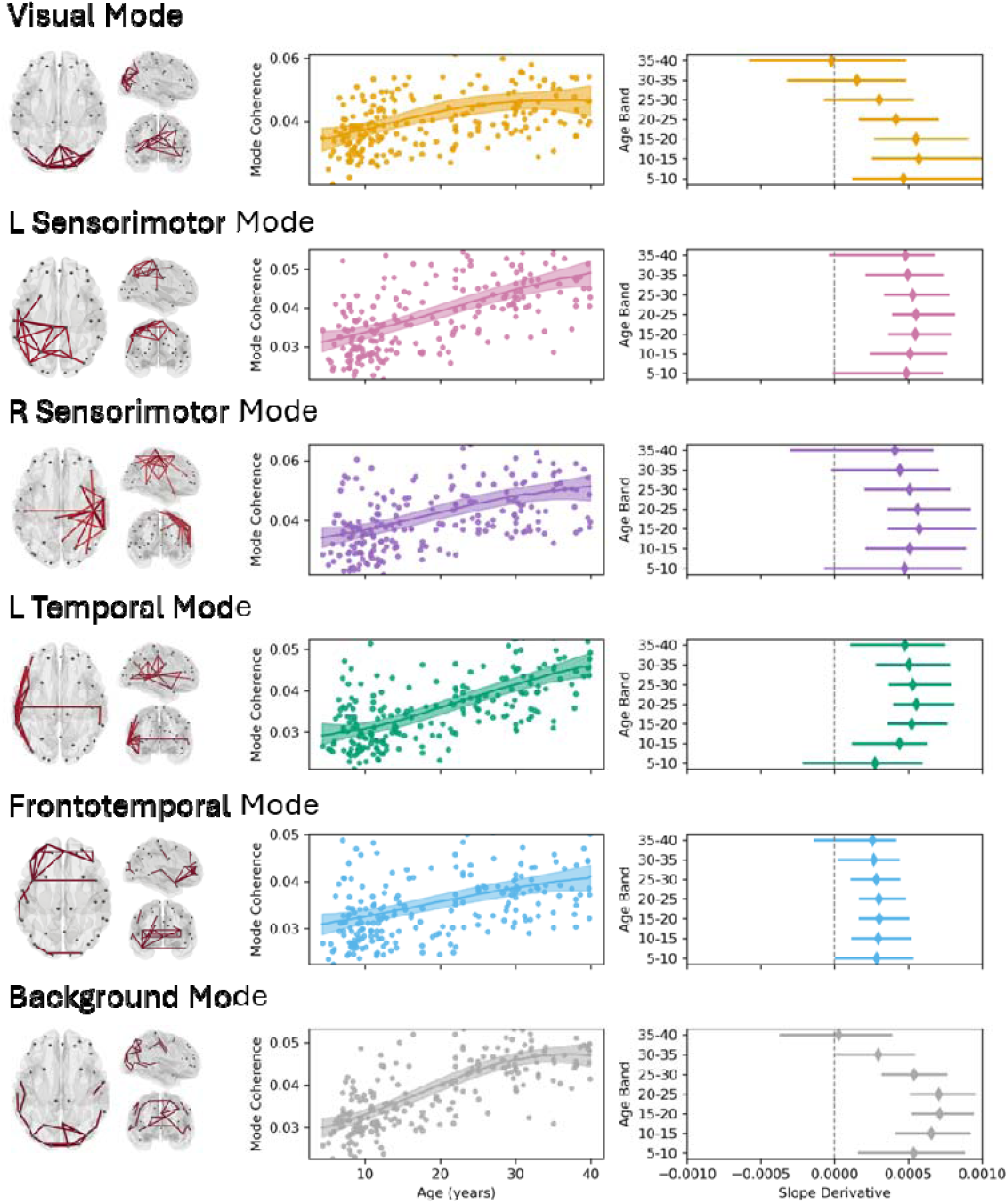
Age-related changes in mode coherence. **Left column:** Lines overlaid on glass brains are the top 5% of edges from the coherence matrix in the theta to beta range (4-30 Hz), at the group level. **Middle column:** Lines are the most probable regression slopes for the relation between age and coherence in the top 5% of edges per mode; shading shows 95% highest density intervals (HDIs). Points are values for individual participants. Outliers above and below the 5^th^ and 95^th^ percentiles, respectively, are not shown. **Right column:** Slope derivatives across discrete age bands. Diamonds denote the most probable slope, bars denote 95% HDIs.

For the benefit of readers who are more comfortable with a traditional (non-Bayesian) statistical framework, we supplemented all our main analyses with corresponding null-hypothesis significance testing (NHST) analyses; results are presented in Supplementary Materials.

## Discussion

In the present study we used DyNeMo, a generative approach for analyzing neural time series data, to characterize developmental trajectories of functional networks underlying emotional face processing. We inferred six whole-brain “modes” elicited during an emotional face processing task; these modes characterized distributed and dynamic functional networks supporting task-related cognition. This work represents a marked departure from previous MEG research in this area which has been constrained to *a priori* regions, frequency bands, and time windows of interest.

Our results provide two key developmental insights. First, different modes inferred by our analyses exhibited distinct developmental trajectories. This was most evident from the results pertaining to dynamic (stimulus-evoked) mode activation levels, where different modes exhibited different directions, rates, and periods of maturation. Here, the visual and frontotemporal modes exhibited particularly notable developmental patterns, with maximal developmental changes occurring during late childhood/adolescence and mid-adulthood respectively. Lateral and dorsomedial prefrontal cortex (both encompassed by the frontotemporal mode) are major components of the canonical face processing network, and have long been associated with emotional processing and broader social cognition, particularly in the context of emotional face processing^28–31^. Nonlinear development of frontotemporal activation may reflect refinement of these cognitive processes. The late visual effect is more surprising, since one would expect the visual system to be fully matured by adulthood. Recent work has provided evidence for protracted, large-scale brain development throughout the lifespan^32^. The observed changes to visual mode activation in early adulthood may reflect late “tuning” of the visual system as part of this ongoing brain maturation. However, further work will be needed to properly characterize this effect. With respect to mode coherence, developmental trajectories were more consistent across modes; in most cases characterized by an approximately linear increase with age, excepting the visual and background modes which showed nonlinear effects. These trajectories are consistent with previous work showing that whole-brain connectivity generally increases with age, even into mid-adulthood^33–37^. Moreover, a few studies have reported similar age-related increases in resting-state connectivity within discrete functional networks, particularly in higher (alpha, beta, gamma) frequency bands^35,37–39^. Our results provide complementary insights into connectivity elicited by an emotional face processing task.

The second major insight from this study is a developmental dissociation between different functional properties (activation and coherence) of the inferred modes. For example, frontotemporal mode activation and coherence exhibited nonlinear and linear relationships with age, respectively. Meanwhile, in the visual mode, age-related changes in activation were only present *after* coherence began to plateau. All of this suggests that mode activation and connectivity are functionally distinct and entail different periods of maturation. Previous work has revealed analogous dissociations between network connectivity and local (i.e., region-specific) task-related responses. For example, the N/M170 face response (typically localized to the fusiform gyri) is generally considered to mature by late adolescence^3,6,40^, while broader connectivity changes associated with face processing can persist into adulthood^34^. In a similar vein, Hunt and colleagues^39^ reported dissociable age-related changes in network connectivity and regional spectral power. Critically, however, our results holistically capture both mode activation and temporally invariant (that is, across the duration of the experiment) connectivity within the same networks, enabling direct comparisons across these two measures. We note that our ability to dissociate network-level *connectivity* from transient network-level *activation* is afforded by DyNeMo’s unique characterization of the latter, which abstracts from multiple task-related neural phenomena including local evoked responses and spectral power changes, as well as transient changes in long-range functional connectivity^23^. The functional dissociations revealed here provide a view which is simultaneously broader in scope (in that it captures multiple discrete processes) and more nuanced than what is typically offered by conventional analyses, which usually characterize “network development” in terms of a single functional property (e.g., ^11,18,21,39^).

With respect to task-related mode activation, it is important to disentangle emotional face processing from more generalized perceptual processing. For example, face presentation might elicit low-level visual processing or preparatory motor activity independently of emotional category. We found that the visual and frontotemporal modes responded differentially to happy and angry faces, indicating explicit involvement in emotional face processing. The latency of these effects (∼600ms post-stimulus in both cases) is noteworthy – while neural discrimination of emotional faces is often detected within the first 100-200ms (e.g., ^3,30,34,41,42^), a few early studies reported effects as late as 1000ms^42,43^; these were broadly attributed to attentional disruptions induced by processing negative emotion. The late frontotemporal discrimination observed here may reflect similar attentional disruption, or perhaps latent evaluative processing (e.g., happy faces are more approachable than angry faces^44^). It is worth noting, however, that evidence for the frontotemporal effect was relatively weak and should be interpreted with caution. Emotional discrimination in the visual mode (for which there was much stronger evidence) may reflect the fact that happy and angry faces differentially affect visual attention^45,46^, which may have latent downstream effects on visual processing; perhaps priming the visual system in anticipation of upcoming stimuli. A final consideration is that the mode time courses are not independent of each other, but instead reflect the relative weighting of mode mixtures through time^22^. Therefore, coincident and opposite activation changes across modes (as we saw here) might be driven by a "true" effect in one mode only. These possibilities warrant further exploration in future studies.

It is also worth addressing age effects in modes that were not sensitive to emotion. Task-related activation in the sensorimotor modes, which declined with age, might reflect sensorimotor dynamics in the absence of a motor response. These dynamics may entail motor-related decision making (neural corelates of which are seen in primary motor cortex bilaterally^47–49^), response inhibition, or a latent effect of motor preparation^50^. However, considering that the age effects were not clearly localized in time relative to stimulus onsets (see S Figure 3), the age effects observed here might simply reflect functional maturation of the sensorimotor system, consistent with previous work in adults^51–53^. The functional role of temporal mode activation is not clear to us. It may relate to response inhibition, since the superior temporal gyrus has been implicated in age-related response inhibition in the context of emotional face processing^28^. However, given the extremely small magnitude of the temporal age effect (near-zero), we are reluctant to draw conclusions about its developmental implications.

This work adds to a small but growing number of studies leveraging DyNeMo to explore resting-state and task-based MEG data^22–24^. The modes inferred here were similar, in terms of their spatial and spectral profile, to those reported in other studies, despite being inferred from different populations and experimental paradigms. Coleman et al.^24^ have suggested that these highly replicable modes reflect a set of intrinsic functional networks, and that perturbation of these networks may capture a range of perceptual and cognitive processes. This perspective will provide a useful common framework for understanding task-related network behaviour.

Here we capitalized on the versatile nature of DyNeMo to capture network-level dynamics underlying emotional face processing in typically developing individuals; our study therefore provides a normative framework on which to build future work in this area. Notably, many neurodevelopmental disorders such as autism spectrum disorder (ASD) are characterized by emotional face processing deficits. In ASD these impairments in social cognition are often accompanied by atypical connectivity profiles in response to faces compared with TDs^11,14–16,18,54,55^, indicative of disrupted network development in this population. Future work could apply DyNeMo to assess how dynamic network behaviour, and associated developmental trajectories, are altered in such disorders. This line of investigation should yield significant insights into atypical brain development as it will capitalize on the broad range of neural phenomena to which DyNeMo is sensitive, as well as benefitting from its highly generalizable and interpretable framework.

## Methods

### Participants

We analyzed 224 MEG datasets from 196 neurotypical individuals (73 female) aged 4.37-39.85 (median age = 15.67). Eighty-nine of these datasets were from the Province of Ontario Neurodevelopmental (POND) Network database; the remaining 135 were archival datasets collected at The Hospital for Sick Children in Toronto. All adult participants and a parent/legal guardian of child participants provided written informed consent according to the Declaration of Helsinki, and children provided verbal assent.

### Experimental Procedures

Participants completed an emotional face processing “Vigilance” task (Figure 6) during MEG recording. Stimuli were images of emotional faces (happy or angry; 26 unique faces, 12 female) from the NimStim Face Stimulus Set^56^ with coloured borders (blue or purple). Each trial began with a face presented for a variable duration of 300-700 ms, followed by a fixation cross for 450-1700 ms. Thus, the inter-stimulus interval (ISI) varied between 750-2400 ms. Participants were instructed to respond with a right-handed button press (using an MEG-compatible button box) to the presence of borders in a target colour, appearing on 25% of trials. Throughout the task, stimulus and fixation cross durations were continuously adjusted to maintain consistent error rates (>=95% for target trials and >=80% for non-target trials) for each participant. Stimuli were presented using Presentation® software (www.neurobs.com).

**Figure 6.**
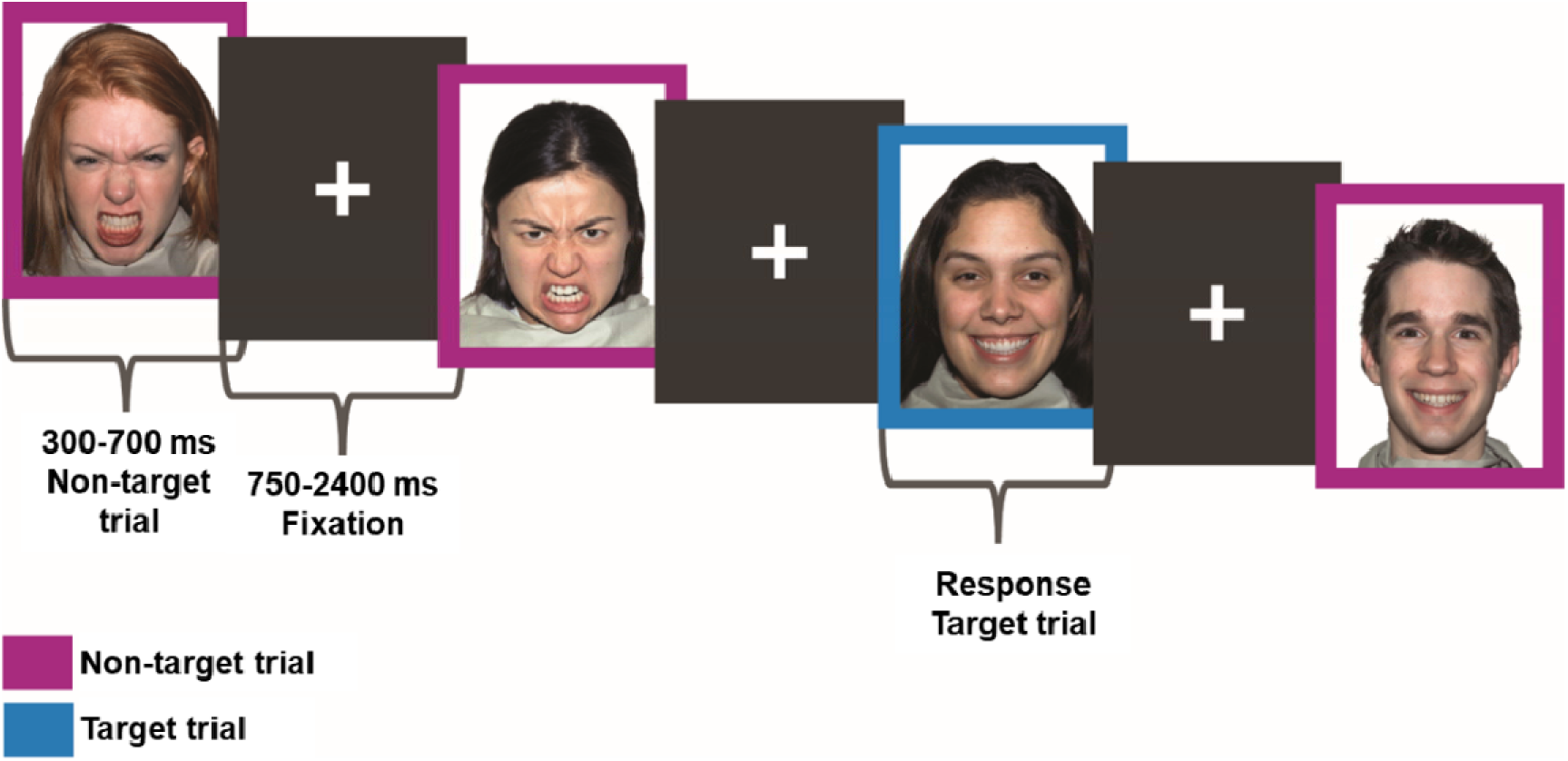
Emotional face processing task. On each trial, participants saw a happy or angry face with a coloured border (blue or purple). Participants were instructed to make a right-handed button press when they saw their target colour border, appearing on 25% of trials.

### MEG and MRI Recording

MEG data were recorded using a 151-channel CTF system (CTF MEG International Services LP, Coquitlam, BC, Canada), housed in a magnetically shielded room. Subjects lay supine and stimuli were presented on an MEG-compatible screen with a viewing distance of 78 cm, subtended ∼ 14 × 16 degrees of visual angle. Recorded data were sampled at 600 Hz with an online 0–150 Hz antialiasing filter, and a third-order spatial gradient applied to reduce environmental noise. Prior to MEG recording, subjects were fitted with fiducial coils (left and right pre-auricular points, nasion) which enabled digitization of anatomical points on the head relative to the sensor array, as well as continuous monitoring of head position during the scan. Following MEG recording, the fiducial coils were replaced with radio-opaque markers, and subjects completed an MRI scan. Structural T1-weighted images were collected using either a Siemens 3.0 T MAGNETOM Trio with a 12 channel head coil (TR = 2,300 ms, TE = 2.96 ms, FOV = 240×256 mm, 192 slices, 1 mm isotropic resolution), or (after a scanner upgrade) a PrismaFIT with a 20 channel head and neck coil (TR = 1,870 ms, TE = 3.14 ms, FA = 9°, FOV = 240×256 mm, 192 slices, 0.8 mm isotropic resolution).

### Data Preprocessing and Beamforming

The following procedures were performed in the Python environment using functions from the MNE library^57^. Raw sensor-level MEG data were downsampled to 250 Hz, band-pass filtered between 1 and 40 Hz, and any channels whose variance exceeded the 75^th^ percentile of variance in all channels by five times the interquartile range were rejected. Artefacts in the filtered, continuous data were annotated in 1 s segments; these included periods of excessive movement (>= 10 mm in any direction from starting head position) and periods in which variance exceeded the 75^th^ percentile by 3 times the interquartile range. Bad segments encompassed a relatively low proportion of timepoints across datasets (mean, max = 2.43%, 18.18%).

For each participant, based on their individual T1 scan, we conducted surface reconstruction and generated a single-shell conduction model^58^ using the boundary-element method with Freesurfer’s *recon-all*^59^ and MNE functions. Raw MEG sensor data were co-registered to individual MRIs using a semi-automated procedure. First, we manually identified coordinates of fiducial points (radio-opaque markers) in the T1 images; these coordinates were transformed to the reconstructed head coordinate frame using the head-to-MRI transformation matrix from recon-all. We then transformed the MEG sensor locations into the same coordinate frame as the head/MRI data. Here, we computed a rigid-body transformation to align the digitized sensor locations (fiducial coil positions) with the MRI fiducials; this alignment was further adjusted using surface-based iterative closest point (ICP) fitting. Co-registration quality was verified by visualizing the aligned sensor positions relative to the reconstructed head surface. Twenty-four datasets had missing or poor-quality corresponding T1 scans; in these cases, we used T1 images from an age- and sex-matched participant for co-registration.

We source-localized each participant’s data to a set of regions throughout the cortical volume. Consistent with recent work^23,24,60^ we derived a set of 38 anatomical parcels from a publicly accessible atlas (https://github.com/OHBA-analysis/osl-ephys/blob/main/osl_ephys/source_recon/files/fmri_d100_parcellation_with_PCC_reduced_2mm.nii.gz) and placed a dipole at the probabilistic maximum of each. Dipole coordinates were then transformed to subject-specific source space, and a forward model calculated for all dipoles. Source-level activity at each dipole was estimated using a linearly-constrained minimum-variance (LCMV) beamformer. We computed data covariance using all available data, excluding bad segments, and regularized at 5% using the Tikhonov method^61^. Dipole orientation was optimised based on the direction of maximum power. Finally, we applied normalization using an estimate of the projected noise to mitigate depth bias introduced by the unit-gain constraint. This provided 38 “virtual electrode” timecourses per subject; we applied symmetric orthogonalization to these timecourses to reduce spurious between-region covariance due to source leakage^62^.

Following beamforming, we applied additional processing steps to prepare the spatially resolved data for generative model training^22,23,63^. We first removed bad segments to prevent model over-fitting to movement and noise, and applied sign flipping to align the polarity of each timecourse across subjects (thus maximizing the between-region covariance structure across subjects). Data were then time-delay embedded (TDE) to insert 14 additional channels (i.e., time-lagged copies of the original channels), with lags of ±7 samples (±30 ms at 250 Hz). TDE enables identification of networks characterized by frequency-specific phase coherence, as well as providing information about the spectral content of the learned modes^64,65^. To avoid computational problems imposed by TDE (which multiplied the number of channels by 15) we reduced the dimensionality of the TDE data using principle component analysis (PCA); we selected 90 principle components as this explained a high percentage (∼74%) of variance in the TDE data. Finally, we standardized (z-scored) the 90-channel time series.

### Dynamic Network Modes

Dynamic Network Modes (DyNeMo)^22^ is a generative modelling approach for characterizing dynamic functional connectivity in neural time series data. The observed data are assumed to arise from a set of latent variables, or spatial *modes*, each defined by multivariate normal distribution known as the *observation model*. The observation model comprises a mean vector and covariance matrix which capture the mode mean (typically fixed at zero) and connectivity structure respectively^22^. Temporal dynamics in mode activation are captured by a weighted mixture of the mode means and covariances at each time point; weightings are determined by a set of mixing coefficients, whose temporal evolution are learned by a recurrent neural network (RNN).

While DyNeMo is mainly data-driven, the number of (to-be-learned) modes must be specified *a priori*. Previous work has shown that six modes are sufficient to capture as many distinct bilateral networks^23,63^; introducing more modes can have the effect of “splitting” modes into (e.g., unilateral) subdivisions^22^. As such, we chose to infer 6 modes in this study. When working with TDE data, modes are typically learned based on data covariances (as opposed to means); we therefore set the mean vector of each mode to zero and did not update its value during training^22,60,65^. An important consideration is that separate training runs on the same data can converge to different latent descriptions of the data (that is, sets of modes with different spatiospectral profiles). Following previous recommendations^60^, we conducted five independent training runs on the same data and selected the model with the lowest variational free energy for subsequent analysis (spatial modes were generally consistent across the 5 runs; see S Figure 7).

After model training and selection, we extracted the learned mixing coefficients and normalized them according to mode covariances^60^, providing a set of positive-definite mode timecourses summing to 1.0 at each time point. We additionally inferred spectral information associated with each mode (change in power spectral density, PSD, and coherence) using the GLM-spectrum method^66^. This method also produces a static PSD corresponding to the intercept term in the GLM (i.e., the mean over timepoints). We then created power maps based on the change in power associated with each mode, compared to the static PSD. Note that the PSDs shown in Figure 1 (right-hand column) were computed as the weighted average across regions for that mode; weightings were proportional to the power change in each region relative to the static PSD.

Mode timecourses were epoched relative to two types of experimental events: (1) button-press responses, and (2) the onset of emotional face stimuli on correct non-target trials. Epochs were defined as -200 to 1000 ms relative to either event. With respect to emotional faces, we selected only correct trials because (inappropriate) manual responses could confound mode activation relevant to face processing.

### Statistical Procedures

We took a fully Bayesian approach to group-level analyses. The following procedures were implemented using functions from the *brms*^67^ and *BayesFactor*^68^ R packages, implemented in the Python environment via the *rpy2* library (https://rpy2.github.io/).

### Time-Domain Analyses of Happy Versus Angry Faces

Each participant was associated with a set of 1200 ms epochs (including 200 ms pre-stimulus), time-locked to the onset of a happy or angry face. We averaged these epochs (separately for happy and angry) within participants while preserving the time dimension; this rendered one 1200 ms timecourse per participant, emotional condition, and mode. At each timepoint, we conducted a Bayesian paired *t*-test between the vector of activation values elicited by happy faces versus that of angry faces (similar to previous time-domain Bayesian analyses^27,69–73)^. This yielded a Bayes factor BF_10_ per timepoint and mode. These BF_10_ values can be interpreted as a continuous measure of quantitative strength of evidence for a true correlation (see ^26,27,74,75^ for introductions to Bayes factors and their interpretation). Bayes factors were computed independently for each mode. Note that BF_10_ values are interpretable at face value and do not require multiple comparison correction^27,75^.

### Analyses of Nonlinear Developmental Trends in Mode Activation and Coherence

Mode activation was defined as the mean over the 0-750ms period (750ms was the minimum ISI) in the epoched mode timecourses for correct non-target trials (collapsed across emotions). Thus, mode activation was represented by a single value per mode, participant and epoch. Mode coherence was defined as each participant’s mean coherence in the 4-30 Hz range over network edges (i.e., cells of the mode coherence matrix) where values were maximal (top 3%) at the group level. Thus, coherence was represented by a single value per participant, per mode.

For each mode we implemented a Bayesian generalized additive model (GAM) which modelled a nonlinear effect of age on the dependent measure (mode activation or coherence) using thin-plate regression splines^76^. When analyzing mode activation, we included a random intercept term for unique scan IDs (to account for multiple epochs per scan). Prior to model fitting, we re-scaled age, mode activation and mode coherence values using independent *Z* transforms. This allowed us to define scale-agnostic (i.e., interpretable) priors and also improved computational efficiency. Parameter estimates for each mode were back-transformed to their original scales for plotting. See Supplemental Materials concerning priors and detailed model fitting parameters.

To better characterize the developmental trajectory of each mode, we defined seven discrete bands across our age spectrum (5-9, 10-14, … 35-40); each band spanned 4 years, with the exception that the final band (35-40) spanned five years so as to capture the full age range of our sample. Within each age band, we estimated the local slope (i.e., steepness) of the smooth age function based on finite-difference estimates. We drew posterior samples from the smooth age function (using *posterior_smooths()* from *brms*) at each one-year increment of age +/- a small finite difference step Δ, and calculated their derivative as:

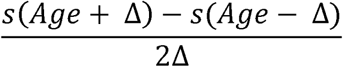

where *s* represents posterior draws from the smooth Age function and Δ = 0.1. This provided a distribution of derivative-based slope estimates for each 1-year age interval. Finally, we averaged these distributions within each band, producing one distribution of slope estimates per age band. We computed 95% highest density intervals (HDIs) from these distributions.

### Rationale for Parameter Estimation

For the developmental analyses we embraced parameter estimation via multilevel Bayesian regression. This is a generative modelling approach which characterises the *posterior distribution* (i.e., joint probability) of parameters over a range of plausible values, given the model and the observed data^77,78^. In the case of a linear regression, for example, we would estimate the joint posterior for the intercept and slope (α, ß respectively) to characterise the relationship between two observed variables (e.g., age and mode amplitude)^‡^. This approach allowed us to robustly quantify the magnitude and credible range of age-related effects in our data. Moreover, the multidimensional nature of the posterior distribution means that the spread of probability on a given parameter can be captured conditionally across a range of some predictor variable. Stated differently, this approach allowed us to robustly quantify the most plausible range of slope values per age band. In turn, we can make quantitatively supported statements about the rate(s) of developmental change within a given age range based on highest-density intervals (HDIs). HDIs are a means of summarizing the posterior distribution that represents the most credible range of values at some level of confidence – here we used 95% HDIs. This allows us to make concrete probabilistic statements about the distribution of the parameter in question. For example, if the HDI for slope includes only positive values, then we can say *with 95% confidence* that the true value of the slope is positive, given the observed data and the model. In this example we may also conclude that zero (which lies outside the HDI) is not credible, with the same level of confidence^77,78^.

## Code Availability Statement

Code for all analyses reported in this manuscript are publicly available on GitHub (https://github.com/lyambailey/Dynamic_Networks_EGNG_TDC).

## Acknowledgements

We are grateful to Jon Fawcett for his advice concerning model fitting with brms, and to Giang Bui for assisting with MEG-MRI co-registration. This research was conducted with the support of the Ontario Brain Institute (POND, PIs: Anagnostou/Lerch), an independent non-profit corporation, funded partially by the Ontario government, Canadian Institutes of Health Research (MOP: 119541) and (PJT-178370); LB was partially supported by a Restracomp Fellowship awarded by The Hospital for Sick Children.

## Supplementary Materials

**S Figure 1.**
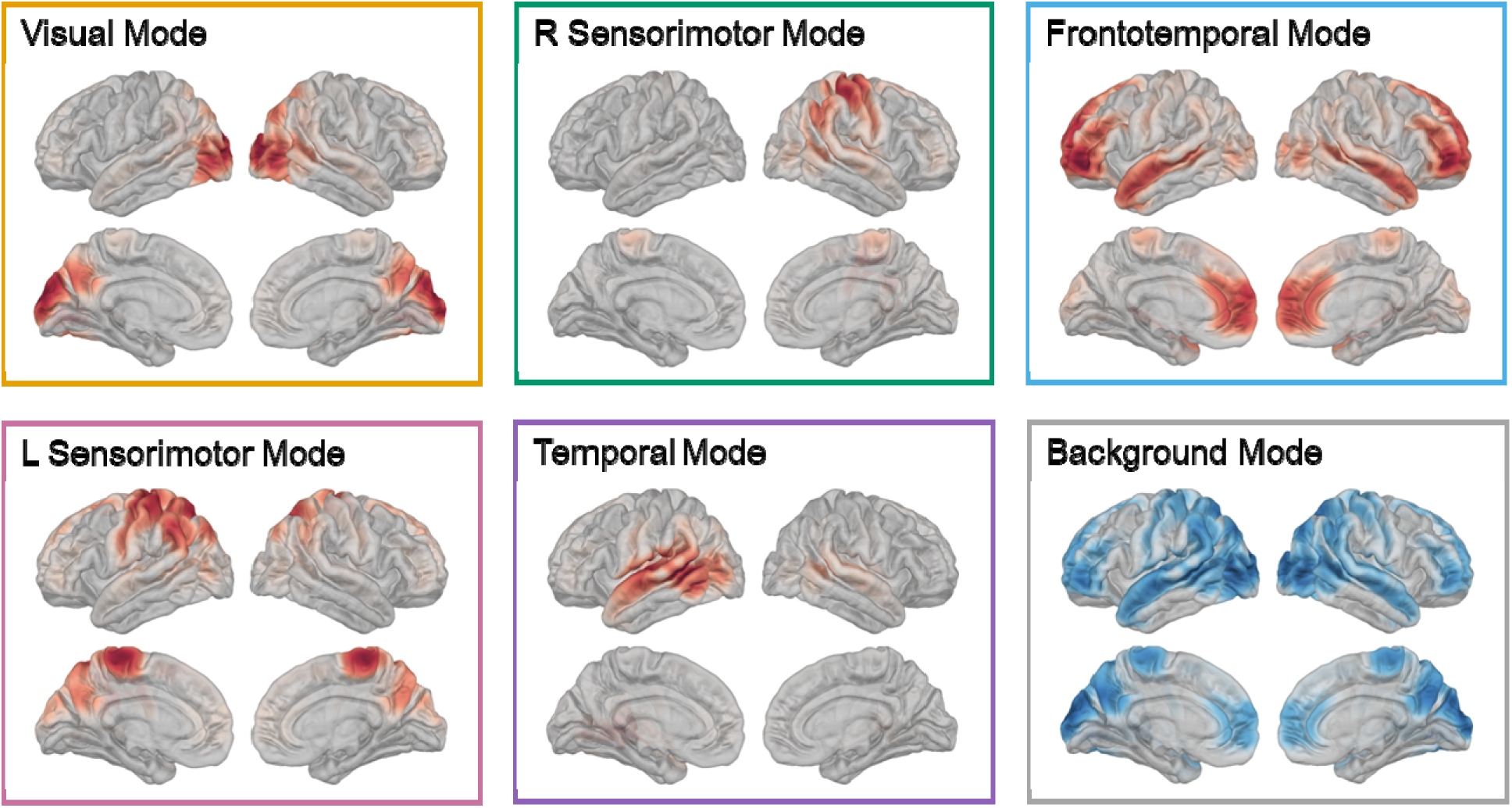
Spatial maps on lateral and medial brain surfaces show power change from the static PSD, for each of the six inferred modes.

**S Figure 2.**
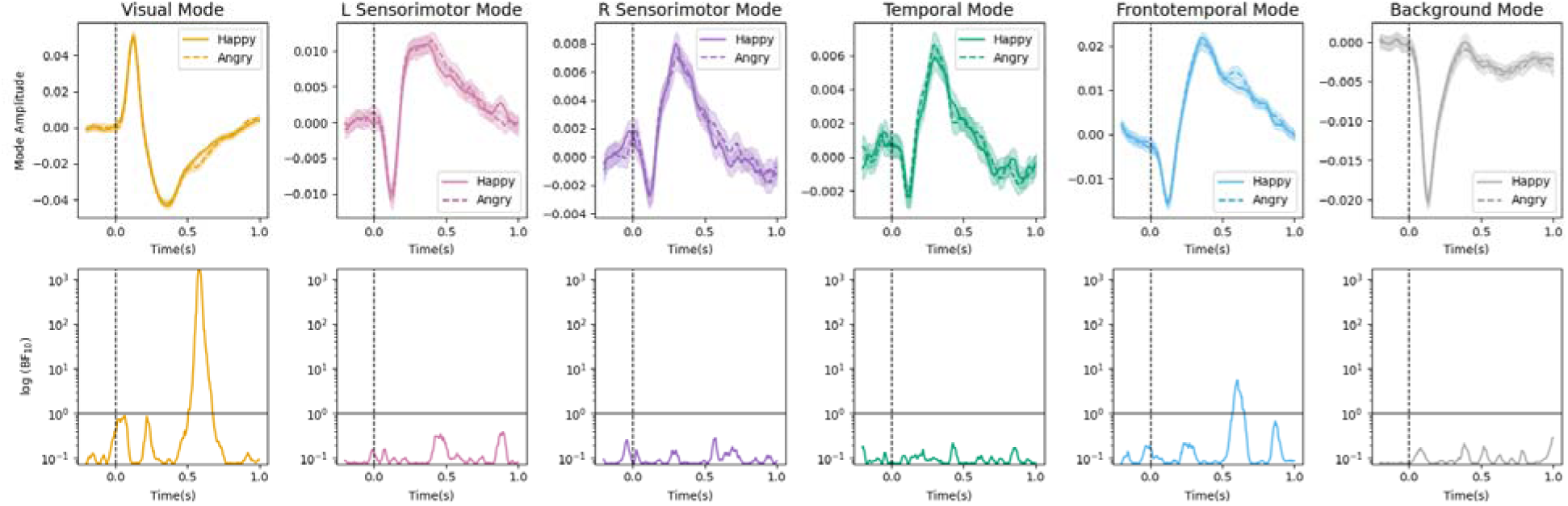
Effect of emotion on mode activation. **Top row:** Mean mode timecourses relative to the onset of happy (solid lines) and angry (dashed lines) faces. Shading reflects ±1 SE. Timecourses were baseline-corrected for visualisation purposes only. **Bottom row:** Log-transformed Bayes factors (BF_10_) computed for each time point, reflecting the strength of evidence for the difference between happy and angry faces. Solid horizontal lines denote BF_10_ = 1.0, above which values support a difference between conditions. Dashed vertical lines in all panels denote time of stimulus onset.

### Time-domain analysis of developmental effects

We investigated which time points following face presentation were driving the observed age effects shown in Figure 4. For each time point in each mode timecourse, we conducted a Bayesian correlation between the corresponding amplitude values (one per subject) and participant ages. This yielded a Pearson correlation value (*r*) and Bayes factor (BF_10_) for each timepoint, per mode. The Bayes factors may be interpreted as a quantitative measure of strength of evidence for a true (anti-) correlation between age and mode amplitude.

**S Figure 3.**
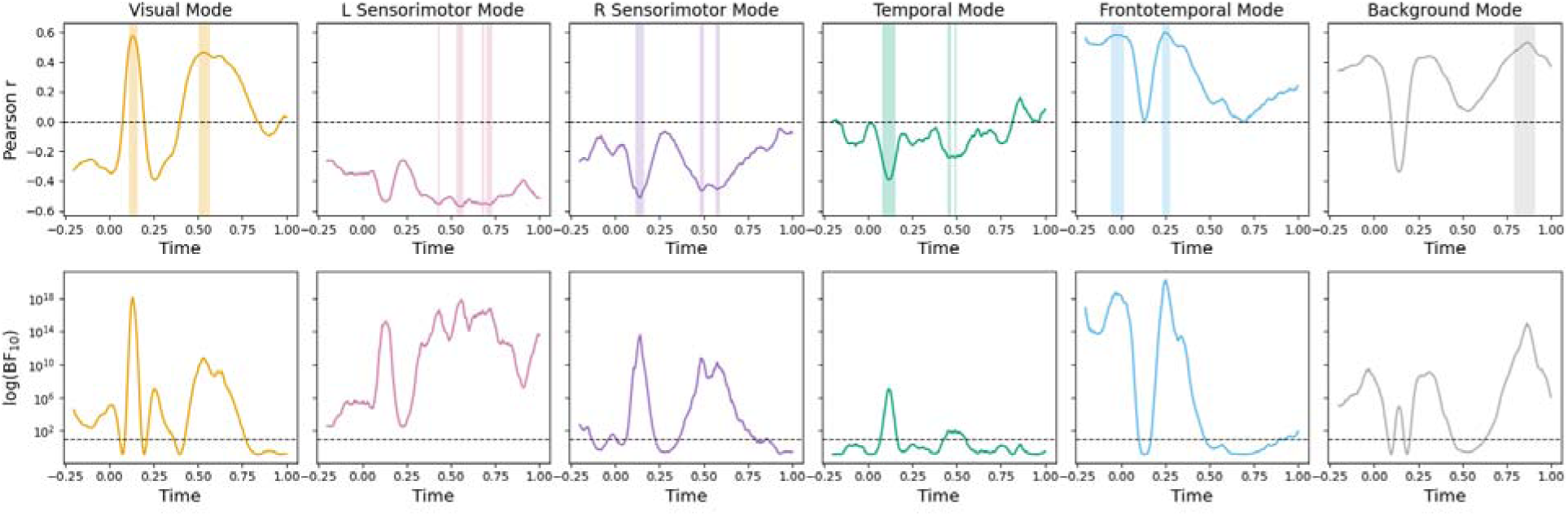
Time-domain analysis of age effects on mode activation. **Top row:** Pearson correlations (*r*) between age and mode amplitude, at each time point in each mode timecourse. Horizontal dashed line indicates *r* = 0. Shaded regions denote periods of highest evidential strength (top 10% of corresponding BF_10_ values). **Bottom row:** Log-transformed Bayes factors (BF_10_) at each time point, representing quantitative strength of evidence for a true (anti)correlation. Horizontal dashed line indicates BF_10_ = 1.0, above which values support a true correlation.

Age-amplitude correlations and Bayes factors in the time dimension are shown in S Figure 3. For the visual and frontotemporal modes, peak positive correlations (and accompanying peaks in strength of evidence) aligned with peaks in the corresponding average mode timecourses (Figure 2; around 100ms and 250ms respectively), suggesting that age predominantly modulated stimulus-related increases in visual and frontotemporal mode activation. Age also modulated, to a lesser extent, the frontotemporal response at stimulus onset, and the period of visual deactivation (∼500 ms). For the temporal and right sensorimotor modes, the strongest evidence for a negative correlation aligned with the visual mode activation peak, during which all other modes experienced some degree of deactivation (see Figure 2). This, alongside the age-related changes in visual deactivation, suggests that the age effects for these modes might be driven by the degree of trade-off between networks. Finally, negative correlations between the left sensorimotor mode and age were of similar magnitude (and strength of evidence) between approximately 500-800 ms, indicating a nonspecific effect of age on this mode. Interestingly, all but the temporal mode exhibited non-zero correlations during the pre-stimulus period, suggesting that age also affected mode activation unrelated to face processing.

### Disaggregating developmental trajectories by sex

For both the mode amplitude and mode coherence analyses, we initially fit separate smooth terms for males and females. As stated in the main text, we did not see credible evidence for sex-based differences in developmental trajectories of mode activation or mode coherence. These results are illustrated in S Figures 4 & 5 respectively. Moreover, model comparisons with leave-one-out cross-validation (implemented using the *loo* R package) indicated a negligible benefit of including sex in any model. Differences in expected log predictive density (ELPD) were small for all modes in both sets of analyses (0.1-2.5); corresponding LOOIC differences (analogous to conventional AIC differences) were similarly negligible (0.2-4.9). Moreover, ELPD differences were similar to or smaller than their corresponding standard errors (se_diff = 0.5-3.0). We concluded that, for all modes in both sets of analyses, the two models (with and without sex disaggregation) had statistically indistinguishable predictive performance.

**S Figure 4.**
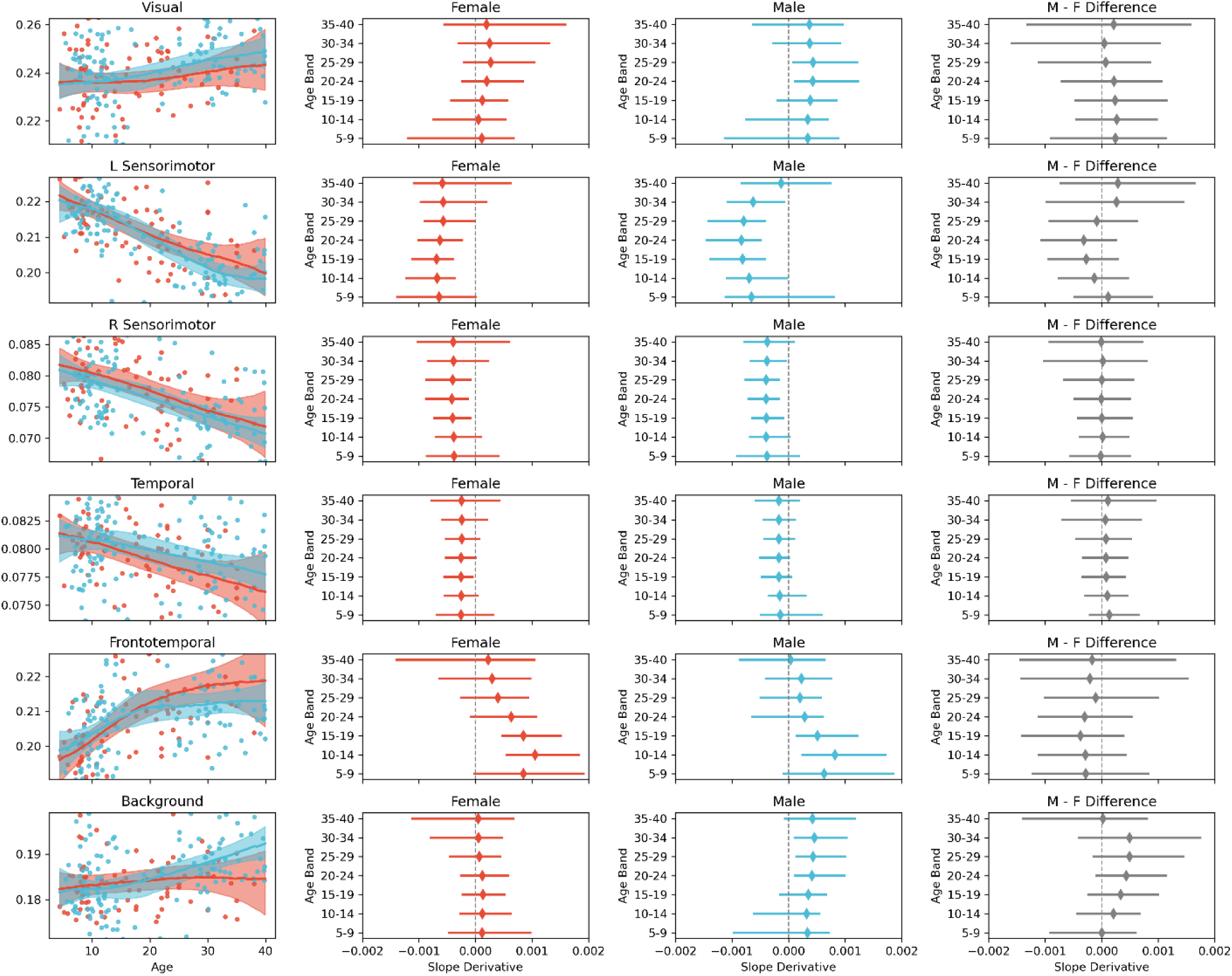
Developmental trajectories in mode amplitudes in the first 750ms following emotional face presentation, disaggregated by sex (blue = males, red = females). **First column (left):** Lines are the most probable regression slopes for the relationship between age and mode amplitude. Shading shows 95% highest density intervals (HDIs). Points are values for individual participants (outliers with values exceeding the 5^th^ and 95^th^ percentile are not shown). **Second and third columns:** Slope derivatives across discrete age bands, separately for males and females. Diamonds denote the most probable slope, bars denote 95% HDIs. **Fourth column (right):** Sex differences in slope derivates per age band, computed by subtracting the posterior distribution of derivates for females from that of males.

**S Figure 5.**
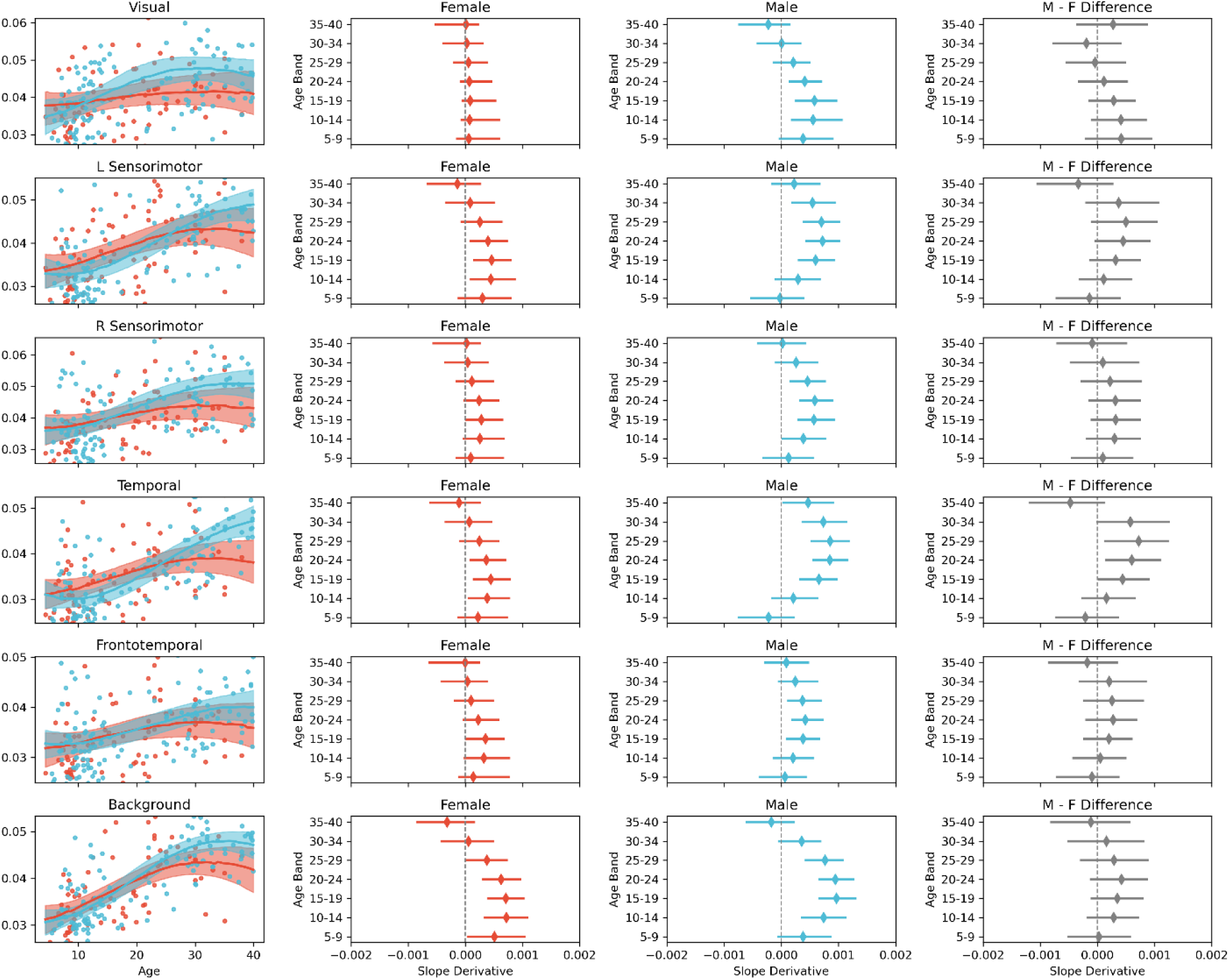
Developmental trajectories in mode coherence, disaggregated by sex (blue = males, red = females). **First column (left):** Lines are the most probable regression slopes for the relationship between age and coherence. Shading shows 95% highest density intervals (HDIs). Points are values for individual participants (outliers with values exceeding the 5^th^ and 95^th^ percentile are not shown). **Second and third columns:** Slope derivatives across discrete age bands, separately for males and females. Diamonds denote the most probable slope, bars denote 95% HDIs. **Fourth column (right):** Sex differences in slope derivates per age band, computed by subtracting the posterior distribution of derivates for females from that of males.

### Null hypothesis significance testing (NHST)

The following describes conventional (non-Bayesian) analyses that are complementary to those described in the main text.

### Time-domain analyses of mode responses to happy vs angry faces

Our time-domain comparison of mode responses to happy and angry faces proceeded as described in the main text; the only difference being that we conducted a regular paired *t*-test (rather than a Bayesian t-test) at each time point. *t*-tests were implemented using the *ttest_rel* function from the S*ciPy* library^1^. This yielded a *p* value for every time point per mode; *p* values were corrected for false discovery rate (FDR) at 1%. The results of this analysis (S Figure 6) revealed significant differences between conditions in the visual and frontotemporal mode, around 550-650 ms following stimulus presentation. This between-condition difference survived FDR correction in the visual mode, but not the frontotemporal mode. This pattern of results is entirely consistent with (and was already evident from) the corresponding Bayes factor analysis in which we found markedly stronger evidence for an effect in the visual mode, and where results can be interpreted at face value without the need for multiple comparison correction^2,3^.

**S Figure 6.**
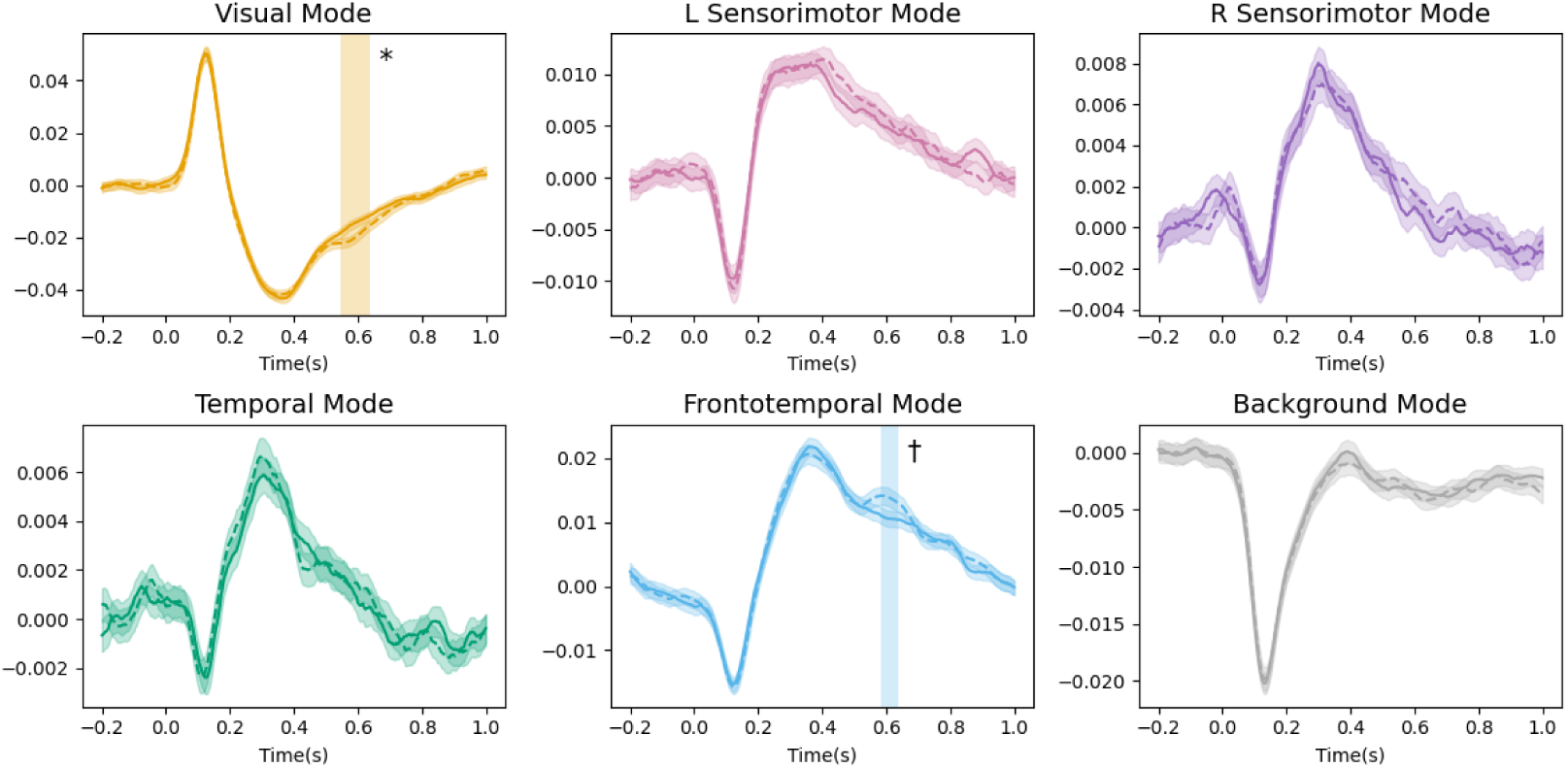
Effect of emotion on mode activation in all modes. In each panel, lines are mean mode timecourses relative to the onset of happy (solid lines) and angry (dashed lines) faces. Shaded ribbons reflect ±1 SE. Timecourses were baseline-corrected for visualization purposes only. Vertical shading indicates periods in which happy and angry timecourses were significantly different. * *p_FDR_* < 0.01; † *p_uncorrected_* < 0.01.

### Nonlinear developmental effects on mode activation and coherence

In the main text we present parameter estimates relating mode activation and coherence to age, obtained with Bayesian generalized additive models (GAMs). We also used non-Bayesian GAMs to test the significance of these nonlinear relationships. GAMs were implemented using the *mgcv* R package^4^, with identical model structures to those described for our Bayesian analysis. As with our Bayesian analysis, all input variables were z-scored prior to analysis. These analyses revealed significant nonlinear relationships between age and mode activation / coherence in all cases; *p* < 0.01 following Bonferroni correction for multiple comparisons. Detailed results for each mode are shown in S Table 1.

**S Table 1.** Results from non-Bayesian analyses of developmental effects. *F* and *p* values correspond to the nonlinear effect of age on mode activation or mode coherence. *p* values are corrected for multiple comparisons (Bonferroni correction).

| Mode | Mode Activation |  | Mode Coherence |  |
| --- | --- | --- | --- | --- |
| | $F$ statistic | $p$ value | $F$ statistic | $p$ value |
| Visual | 6.85 | $< 1.2 \times 10^{-15}$ | 14.76 | $< 1.2 \times 10^{-15}$ |
| L Sensorimotor | 54.23 | $< 1.2 \times 10^{-15}$ | 101.20 | $< 1.2 \times 10^{-15}$ |
| R Sensorimotor | 72.99 | $< 1.2 \times 10^{-15}$ | 21.87 | $< 1.2 \times 10^{-15}$ |
| Temporal | 23.99 | $6.66 \times 10^{-6}$ | 131.50 | $< 1.2 \times 10^{-15}$ |
| Frontotemporal | 12.15 | $< 1.2 \times 10^{-15}$ | 39.62 | $< 1.2 \times 10^{-15}$ |
| Background | 20.22 | $2.60 \times 10^{-5}$ | 64.88 | $< 1.2 \times 10^{-15}$ |

### Results of independent DyNeMo training runs

As stated in the main text, we trained DyNeMo on MEG data across five independent training runs, and selected the run with the lowest variational free energy for further analysis. Spatial power maps for the five runs are shown in S Figure 6.

**S Figure 7.**
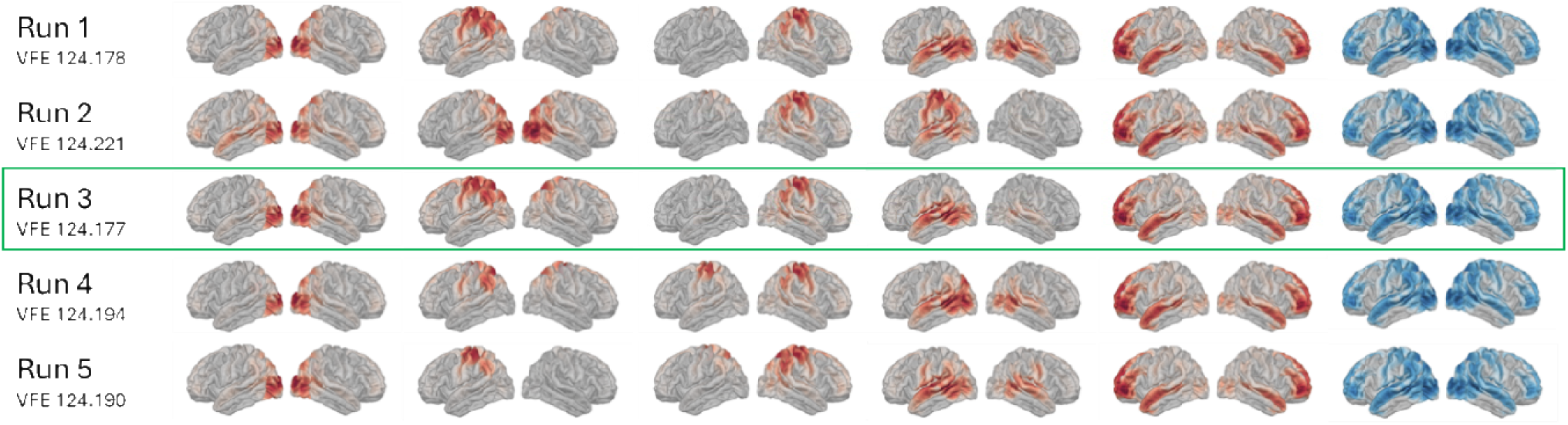
Spatial modes across five independent DyNeMo training runs. The green outline indicates the run with the lowest variational free energy (VFE) selected for further analysis in this study.

### Bayesian priors and model fitting parameters

For all Bayesian generalized additive models (implemented with *brms*), we defined weakly informative priors for the intercept and linear component of the smooth age term: Normal(0,1). For the residual error standard deviation and (when applicable) the standard deviation of subject-level random effects, we used half-Normal(0,1). These priors did not reflect strong predictions about the data but provided better regularization and more reasonable predictions compared to brms’s default (uniform) priors, since they placed higher probability density on parameter estimates close to zero. We defined a more conservative prior for the smooth-term standard deviation, which regularized nonlinear age effects: half-Normal(0, 0.25). This prior captures gentle, global nonlinear trends over the full age range while discouraging sharp, localized fluctuations. This conservative regularization was particularly important in our analyses because our sample was not uniformly distributed across the full age range, meaning that sparsely sampled ages might be prone to spurious nonlinearity. All models were fitted with 4 chains, 20,000 iterations, and 5,000 warmup samples. All other *brms* settings were default. Model convergence was verified using the standard metric of R-hat ≈ 1.0. For the coherence analyses there were zero divergent transitions for any model; for the mode activation analyses, we saw a negligible number of divergent transitions (max <0.05%) in the temporal and right sensorimotor modes, and zero divergent transitions in other modes

## Footnotes

* For reference, BF_10_ values between 3.0 and 10.0 are generally interpreted as “moderate” evidence against the null hypothesis; values > 10.0 are interpreted as “strong evidence” ^25–27^.

† See supplementary materials for results of equivalent NHST.

‡ We give the example of linear regression because these parameters will be intuitive and familiar to most readers. In spline-based nonlinear regression, used in this study, we parameterize the intercept and standard deviation of the spline basis coefficients which determine the local curvature (i.e., “wiggliness”) of the smooth term.

## Notes

https://github.com/lyambailey/Dynamic_Networks_EGNG_TDC

### Competing Interest Statement

The authors have declared no competing interest.

