## Supplementary Materials for "Charting developmental trajectories of dynamic brain networks during emotional face processing"

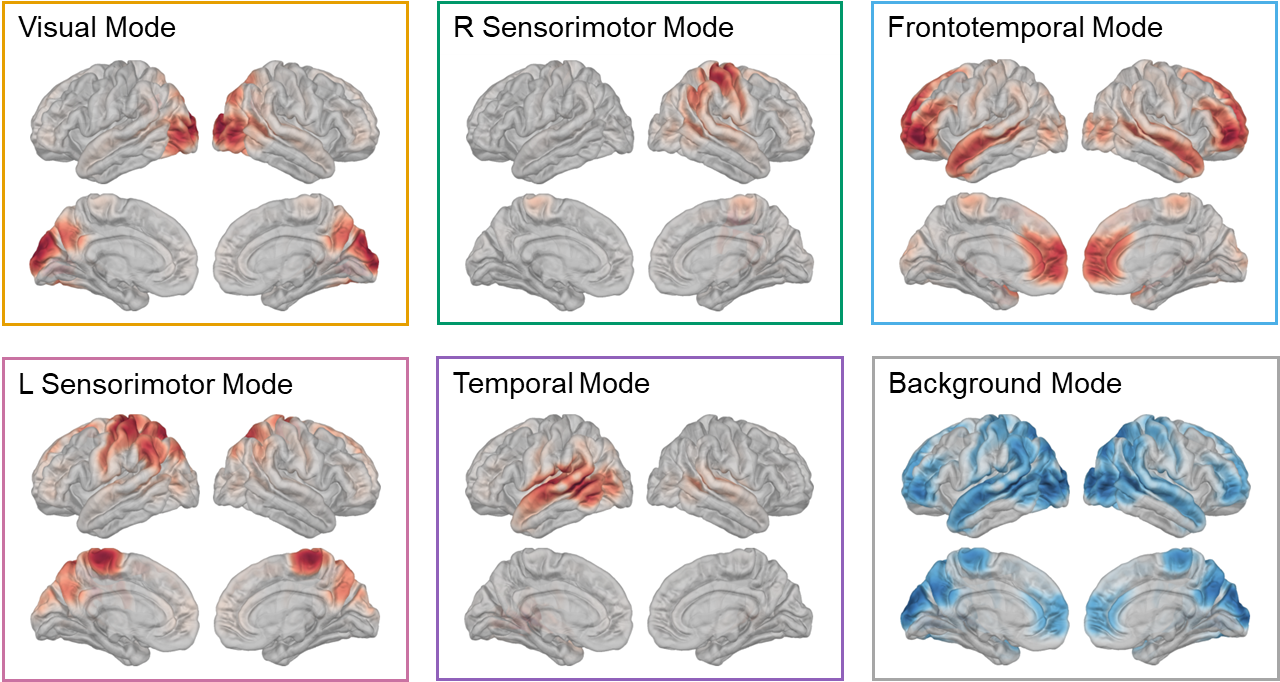

**S Figure 1.** Spatial maps on lateral and medial brain surfaces show power change from the static PSD, for each of the six inferred modes.

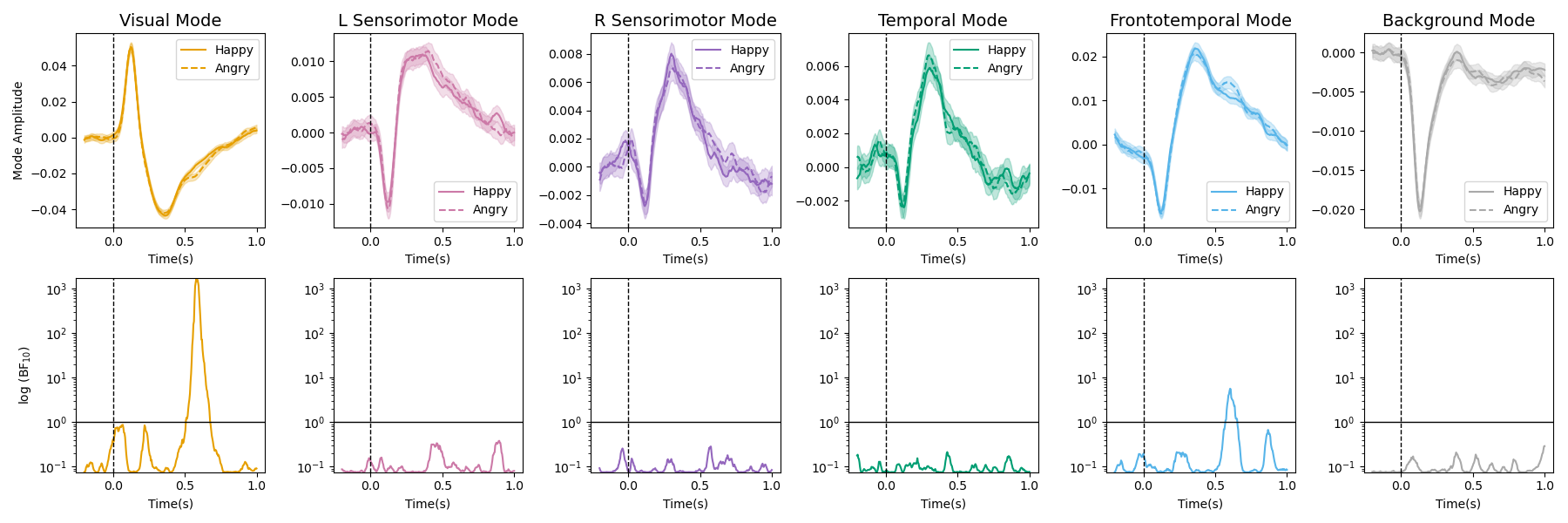

**S Figure 2.** Effect of emotion on mode activation. **Top row:** Mean mode timecourses relative to the onset of happy (solid lines) and angry (dashed lines) faces. Shading reflects ±1 SE. Timecourses were baseline-corrected for visualisation purposes only. **Bottom row:** Log-transformed Bayes factors (BF_10_) computed for each time point, reflecting the strength of evidence for the difference between happy and angry faces. Solid horizontal lines denote BF_10_ = 1.0, above which values support a difference between conditions. Dashed vertical lines in all panels denote time of stimulus onset.

**Time-domain analysis of developmental effects**

We investigated which time points following face presentation were driving the observed age effects shown in Figure 4. For each time point in each mode timecourse, we conducted a Bayesian correlation between the corresponding amplitude values (one per subject) and participant ages. This yielded a Pearson correlation value (*r*) and Bayes factor (BF_10_) for each timepoint, per mode. The Bayes factors may be interpreted as a quantitative measure of strength of evidence for a true (anti-) correlation between age and mode amplitude.

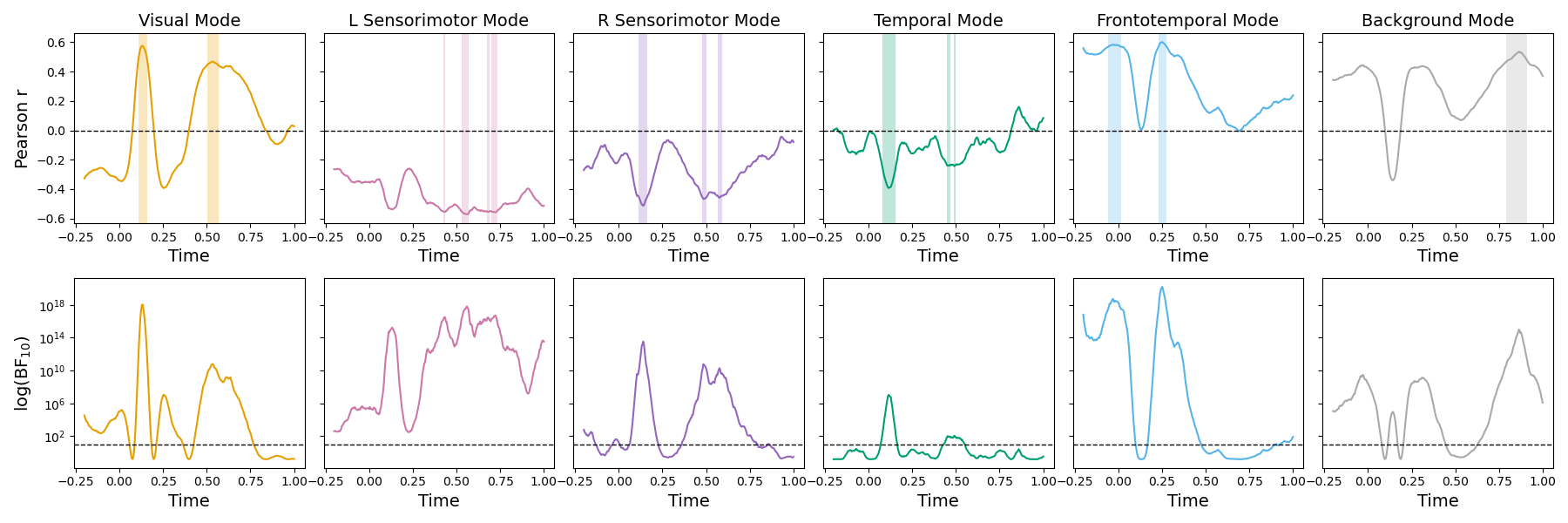

**S Figure 3.** Time-domain analysis of age effects on mode activation. **Top row:** Pearson correlations (*r*) between age and mode amplitude, at each time point in each mode timecourse. Horizontal dashed line indicates *r* = 0. Shaded regions denote periods of highest evidential strength (top 10% of corresponding BF_10_ values). **Bottom row:** Log-transformed Bayes factors (BF_10_) at each time point, representing quantitative strength of evidence for a true (anti)correlation. Horizontal dashed line indicates BF_10_ = 1.0, above which values support a true correlation.

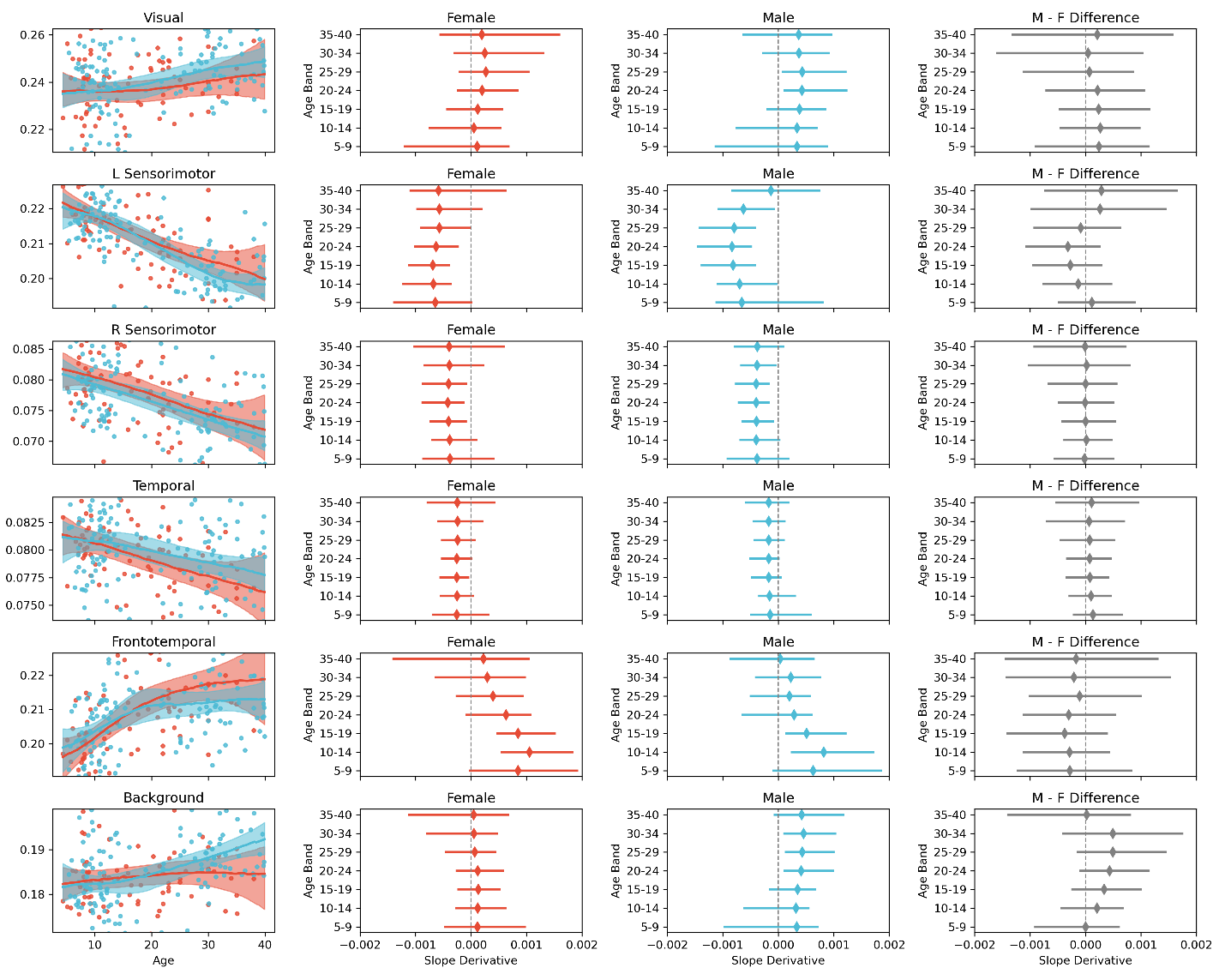

**S Figure 4.** Developmental trajectories in mode amplitudes in the first 750ms following emotional face presentation, disaggregated by sex (blue = males, red = females). **First column (left):** Lines are the most probable regression slopes for the relationship between age and mode amplitude. Shading shows 95% highest density intervals (HDIs). Points are values for individual participants (outliers with values exceeding the 5^th^ and 95^th^ percentile are not shown). **Second and third columns:** Slope derivatives across discrete age bands, separately for males and females. Diamonds denote the most probable slope, bars denote 95% HDIs. **Fourth column (right):** Sex differences in slope derivates per age band, computed by subtracting the posterior distribution of derivates for females from that of males.

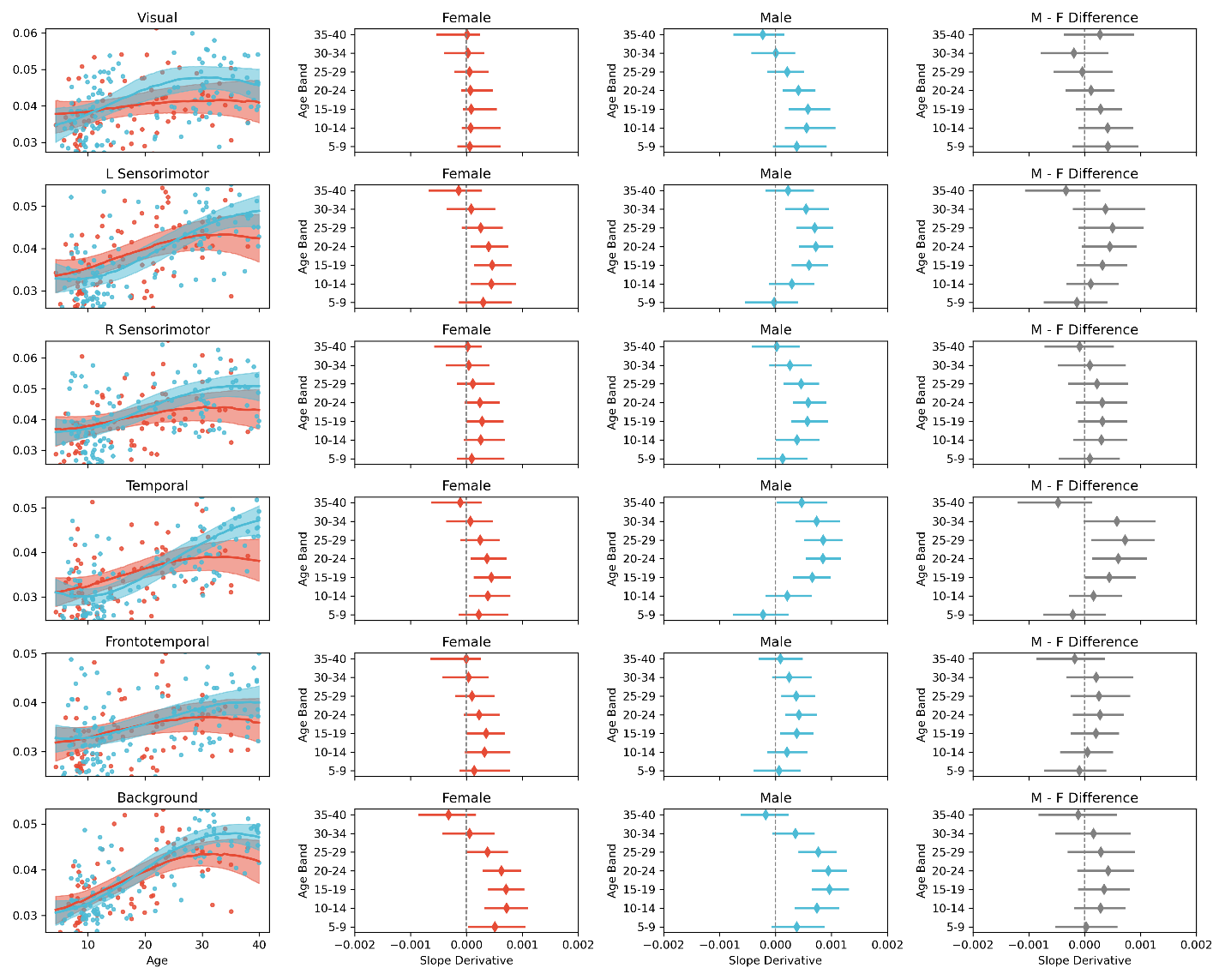

**S Figure 5.** Developmental trajectories in mode coherence, disaggregated by sex (blue = males, red = females). **First column (left):** Lines are the most probable regression slopes for the relationship between age and coherence. Shading shows 95% highest density intervals (HDIs). Points are values for individual participants (outliers with values exceeding the 5^th^ and 95^th^ percentile are not shown). **Second and third columns:** Slope derivatives across discrete age bands, separately for males and females. Diamonds denote the most probable slope, bars denote 95% HDIs. **Fourth column (right):** Sex differences in slope derivates per age band, computed by subtracting the posterior distribution of derivates for females from that of males.

**Null hypothesis significance testing (NHST)**

The following describes conventional (non-Bayesian) analyses that are complementary to those described in the main text.

***Time-domain analyses of mode responses to happy vs angry faces***

Our time-domain comparison of mode responses to happy and angry faces proceeded as described in the main text; the only difference being that we conducted a regular paired *t*-test (rather than a Bayesian t-test) at each time point. *t*-tests were implemented using the *ttest_rel* function from the S*ciPy* library^1^. This yielded a *p* value for every time point per mode; *p* values were corrected for false discovery rate (FDR) at 1%. The results of this analysis (S Figure 6) revealed significant differences between conditions in the visual and frontotemporal mode, around 550-650 ms following stimulus presentation. This between-condition difference survived FDR correction in the visual mode, but not the frontotemporal mode. This pattern of results is entirely consistent with (and was already evident from) the corresponding Bayes factor analysis in which we found markedly stronger evidence for an effect in the visual mode, and where results can be interpreted at face value without the need for multiple comparison correction^2,3^.

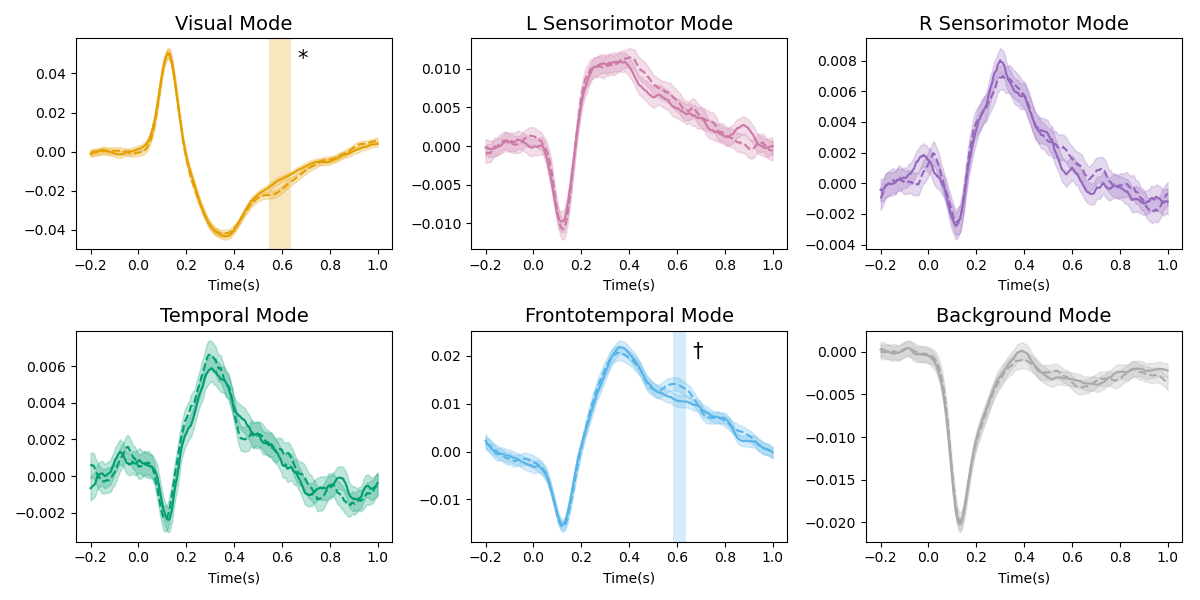

**S Figure 6.** Effect of emotion on mode activation in all modes. In each panel, lines are mean mode timecourses relative to the onset of happy (solid lines) and angry (dashed lines) faces. Shaded ribbons reflect ±1 SE. Timecourses were baseline-corrected for visualization purposes only. Vertical shading indicates periods in which happy and angry timecourses were significantly different. * *p_FDR_* < 0.01; † *p_uncorrected_* < 0.01.

***Nonlinear developmental effects on mode activation and coherence***

In the main text we present parameter estimates relating mode activation and coherence to age, obtained with Bayesian generalized additive models (GAMs). We also used non-Bayesian GAMs to test the significance of these nonlinear relationships. GAMs were implemented using the *mgcv* R package^4^, with identical model structures to those described for our Bayesian analysis. As with our Bayesian analysis, all input variables were z-scored prior to analysis. These analyses revealed significant nonlinear relationships between age and mode activation / coherence in all cases; *p* < 0.01 following Bonferroni correction for multiple comparisons. Detailed results for each mode are shown in S Tables 1 and 2.

**S Table 1.** Results from non-Bayesian analyses of developmental effects. *F* and *p* values correspond to the nonlinear effect of age on mode activation or mode coherence. *p* values are corrected for multiple comparisons (Bonferroni correction).

|  | Mode Activation | | Mode Coherence | |
| --- | --- | --- | --- | --- |
| Mode | *F* statistic | *p* value | *F* statistic | *p* value |
| Visual | 6.85 | < 1.2 x 10^-15^ | 14.76 | < 1.2 x 10^-15^ |
| L Sensorimotor | 54.23 | < 1.2 x 10^-15^ | 101.20 | < 1.2 x 10^-15^ |
| R Sensorimotor | 72.99 | < 1.2 x 10^-15^ | 21.87 | < 1.2 x 10^-15^ |
| Temporal | 23.99 | 6.66 x 10^-6^ | 131.50 | < 1.2 x 10^-15^ |
| Frontotemporal | 12.15 | < 1.2 x 10^-15^ | 39.62 | < 1.2 x 10^-15^ |
| Background | 20.22 | 2.60 x 10^-5^ | 64.88 | < 1.2 x 10^-15^ |

**Results of independent DyNeMo training runs**

As stated in the main text, we trained DyNeMo on MEG data across five independent training runs, and selected the run with the lowest variational free energy for further analysis. Spatial power maps for the five runs are shown in S Figure 6.

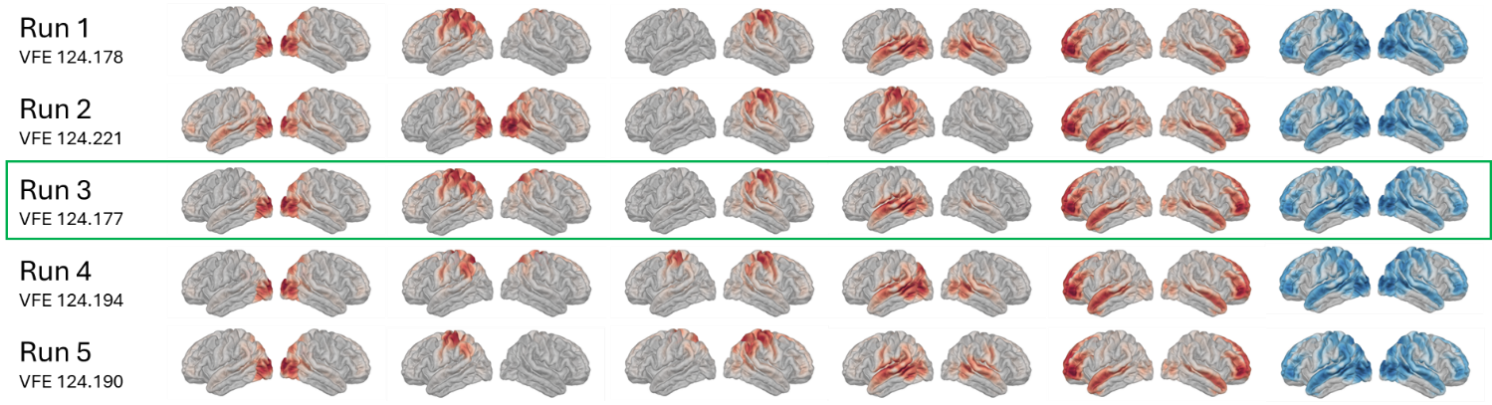

**S Figure 7.** Spatial modes across five independent DyNeMo training runs. The green outline indicates the run with the lowest variational free energy (VFE) selected for further analysis in this study.

**Bayesian priors and model fitting parameters**

For all Bayesian generalized additive models (implemented with *brms*), we defined weakly informative priors for the intercept and linear component of the smooth age term: Normal(0,1). For the residual error standard deviation and (when applicable) the standard deviation of subject-level random effects, we used half-Normal(0,1). These priors did not reflect strong predictions about the data but provided better regularization and more reasonable predictions compared to brms’s default (uniform) priors, since they placed higher probability density on parameter estimates close to zero. We defined a more conservative prior for the smooth-term standard deviation, which regularized nonlinear age effects: half-Normal(0, 0.25). This prior captures gentle, global nonlinear trends over the full age range while discouraging sharp, localized fluctuations. This conservative regularization was particularly important in our analyses because our sample was not uniformly distributed across the full age range, meaning that sparsely sampled ages might be prone to spurious nonlinearity. All models were fitted with 4 chains, 20,000 iterations, and 5,000 warmup samples. All other *brms* settings were default. Model convergence was verified using the standard metric of R-hat ≈ 1.0. For the coherence analyses there were zero divergent transitions for any model; for the mode activation analyses, we saw a negligible number of divergent transitions (max <0.05%) in the temporal and right sensorimotor modes, and zero divergent transitions in other modes
